# Epigenetic Context Defines the Transcriptional Activity of Canonical and Noncanonical NF-κB Signaling in Pancreatic Cancer

**DOI:** 10.1101/2025.06.25.661561

**Authors:** Joana E. Aggrey-Fynn, Joshua Busch, Dominik Saul, Ashish Rajput, Kerstin Willecke, Nicole Klimt, Kothai Rajendran, Nadine Schacherer, Wanwan Ge, Julia Thiel, Amro Abdelrahman, Mark J. Truty, Meng Dong, Steven A. Johnsen

## Abstract

NF-κB signaling can be subdivided into canonical and noncanonical pathways, culminating in the transcriptional activity of RELA and RELB, respectively. However, the upstream signals that activate these transcription factors and their specific regulatory roles in pancreatic ductal adenocarcinoma (PDAC) remain incompletely understood. Here, we demonstrate that TNFα is the primary activator of canonical NF-κB signaling via RELA, while TWEAK engages noncanonical signaling through RELB in PDAC. Using transcriptome-wide gene expression, genome-wide occupancy profiling, and epigenome mapping, we delineate distinct temporal dynamics and regulatory activities of each pathway. Single-cell RNA-seq analysis and multiplex immunofluorescence staining in primary PDAC samples further reveal extensive and distinct interactions with components of the tumor microenvironment. Motif analysis of RELA- and RELB-bound regions uncovers a particularly strong association of RELB with AP1 elements. Notably, while RELA binds to both open and closed chromatin, RELB exclusively occupies regions with pre-existing chromatin accessibility and AP1 co-binding. Collectively, these findings underscore the distinct and complementary roles of TNFα and TWEAK in PDAC, with TNFα engaging a broader transcriptional program via RELA and TWEAK selectively targeting genes marked by accessible chromatin and epigenetic activity. This study provides critical insights into the regulatory dynamics of NF-κB signaling in PDAC, highlighting unique and complementary functions of RELA and RELB in modulating downstream gene expression.

**Significance:** TNFα and TWEAK selectively activate RELA and RELB in pancreatic cancer, revealing that RELB-driven noncanonical signaling depends on chromatin accessibility and AP-1, uncovering distinct, targetable epigenetic dependencies from canonical signaling.

**Graphical abstract:** 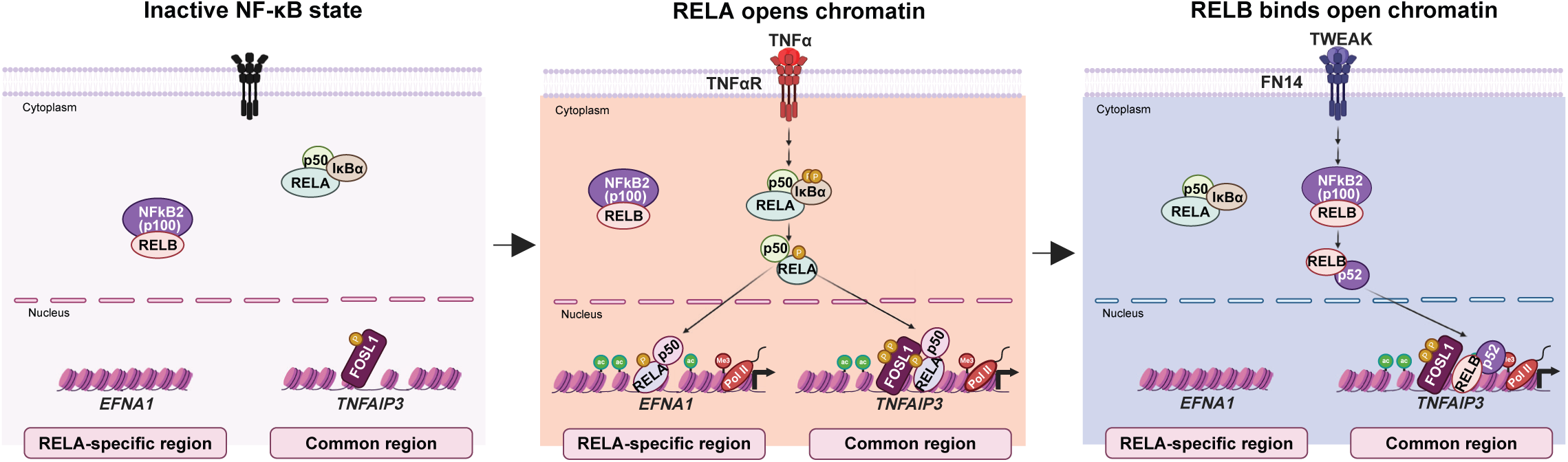

## Introduction

The nuclear factor kappa-light-chain-enhancer of activated B cells (NF-κB) signaling pathway is a pivotal regulator of diverse cellular processes, contributing to cancer development and progression. Activation of NF-κB is frequently observed in pancreatic ductal adenocarcinoma (PDAC), where it is associated with aggressive disease. Studies from our lab and others have linked NF-κB activation to oncogenic KRAS signaling and chronic inflammation, driving enhanced survival signaling, tumor growth, metastasis, and therapy resistance^1–5^. Over 70% of PDAC tumors exhibit aberrant NF-κB activity, correlating with poor prognosis, underscoring its role in regulating key aspects of PDAC progression^1,6,7^.

The NF-κB family comprises several members that form distinct dimers and can be divided into two separate signaling pathways referred to as canonical and noncanonical^8–10^. The canonical pathway is activated rapidly by external stimuli, such as the pro-inflammatory cytokine TNFα, through phosphorylation and activation of the IκB kinases (IKKs)^11,12^. This leads to proteasomal degradation of inhibitory IκB proteins and nuclear translocation of RELA/p50 dimers. The resulting activation is rapid and transient as it simultaneously triggers the expression of negative regulators, including IκBα, A20, and p105, which establish a negative feedback loop^13–15^. In contrast, the noncanonical pathway is slower, relying on the stabilization of NF-κB-inducing kinase (NIK) following degradation of TNFR-associated factor 3 (TRAF3). This allows for the proteasome-dependent proteolytic processing of p100 to p52, releasing RELB/p52 dimers for transcriptional activation^16,17^ (Fig. 1). While the canonical pathway drives inflammation, immunity, and cell survival, the noncanonical pathway has been implicated in lymphoid development, organogenesis, and sustained immune regulation^9–11,13,18^.

**Figure 1:**
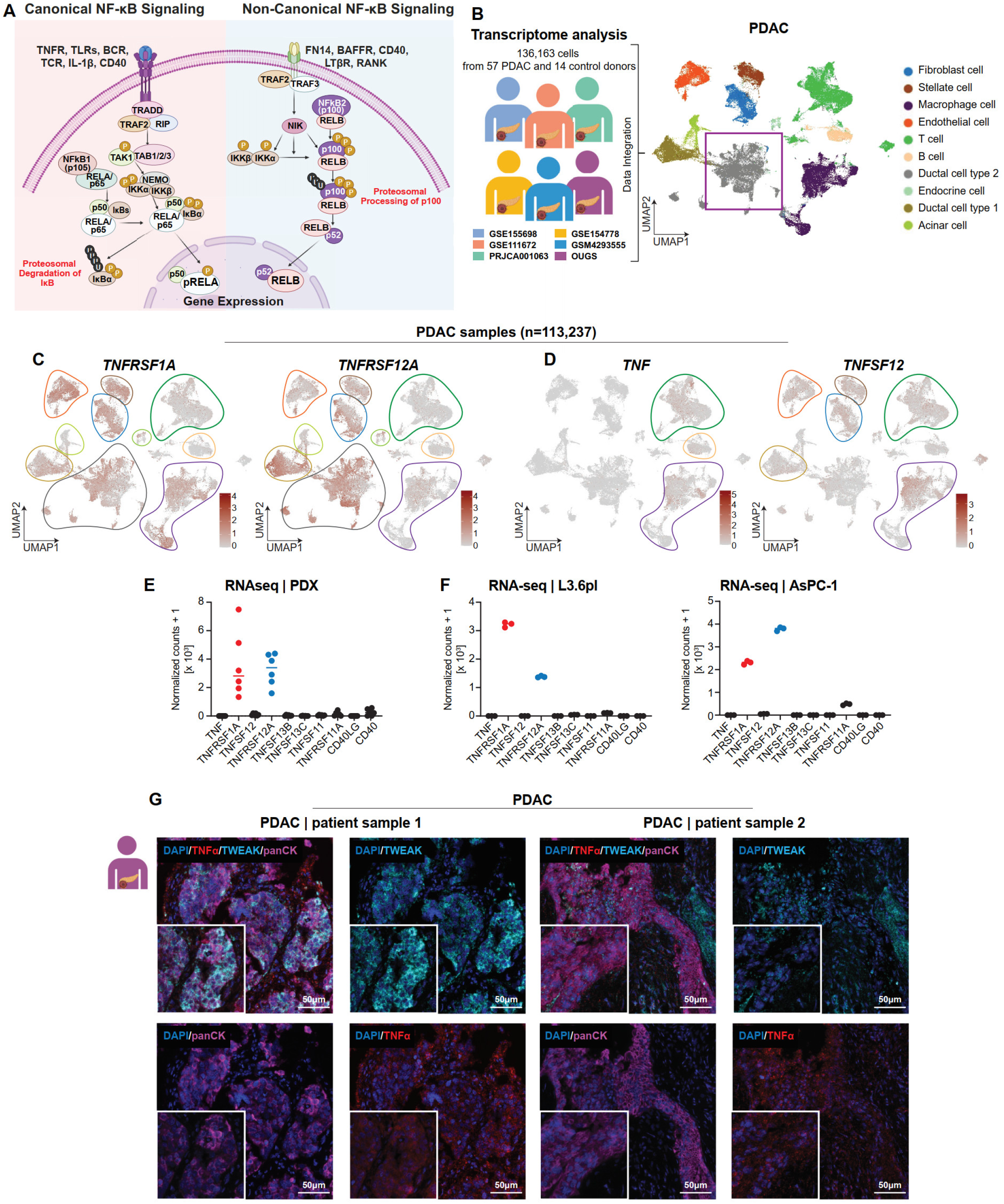
TNFα receptor (*TNFRSF1A*) and TWEAK receptor (*TNFRSF12A*) are the most expressed NF-κB receptors in PDAC. **A**, Schematic representation of canonical and noncanonical NF-κB signaling pathways, illustrating their respective ligand-receptor interactions and downstream activation mechanisms. **B**, Overview of published scRNA-seq datasets (left) and uniform manifold approximation and projection (UMAP) (*right*) of six scRNA-seq datasets from PDAC patients (*n = 136,163 cells, n = 57 donors*, *n = 14 healthy controls*). Cells are colored according to their respective subsets, depicting the cellular landscape of PDAC. **C & D**, UMAP plots showing the expression patterns of TNFαR (*TNFRSF1A*) and TWEAKR (*TNFRSF12A*) alongside their respective ligands TNFα (*TNF*) and TWEAK (*TNFSF12*), highlighting their distinct distribution across cell populations. **E & F**, Bar graphs displaying normalized count values for NF-κB pathway receptors and ligands, derived from RNA-seq data from patient-derived xenografts (PDXs) and PDAC cell lines, demonstrating TNFαR and TWEAKR as the most highly expressed NF-κB receptors in these models. **G**, Multiplex immunofluorescence staining of PDAC tumor samples showing TNFα (red), TWEAK (cyan), and cytokeratin (pan) (purple), providing spatial localization of NF-κB ligands within the tumor microenvironment. Scale bar represents 50 μm (inserts: 20 μm)

Although the canonical pathway has been extensively studied, less is known about noncanonical NF-κB signaling and the differential activities of the canonical and noncanonical pathways remains largely unknown. Moreover, the distinct roles of these pathways in PDAC are poorly understood. Based on current literature, canonical and noncanonical pathways likely have complementary but distinct roles in PDAC biology. Thus, understanding these pathways could uncover new therapeutic targets for PDAC treatment.

Using PDAC patient samples and cell lines, we identified tumor necrosis factor-alpha (TNFα) and its receptor (TNFR), and the TNF-like weak inducer of apoptosis (TWEAK) and its receptor (TWEAKR) as key upstream ligand-receptor pairs potentially driving NF-κB signaling in PDAC. Notably, we demonstrate that TNFα-induced RELA is capable of binding both open and closed chromatin regions and inducing H3K27ac at enhancer regions to activate transcription. Conversely, TWEAK-induced RELB binds exclusively to open chromatin regions, relying on other transcription factors such as AP-1 to activate transcription. These findings were validated through multi-omics approaches, including single-cell RNA-seq, mRNA-seq, ChIP-seq, and ATAC-seq, revealing functional and epigenetic distinctions between the canonical and noncanonical pathways.

This work provides novel insights into the regulatory dynamics of NF-κB signaling in PDAC, highlighting pathway-specific differences. These findings could pave the way for approaches to therapeutic interventions targeting canonical and noncanonical NF-κB pathways to improve outcomes in PDAC.

## Results

### TNF**α**-TNFR and TWEAK-TWEAKR are the most prominent NF-**κ**B signaling pathways active in PDAC

The canonical and noncanonical NF-κB signaling pathways are driven by distinct ligands, receptors, and downstream signaling cascades (Fig. 1a). To identify the major NF-κB signaling pathways active in PDAC, we integrated and analyzed six publicly available single-cell RNA-seq datasets encompassing 57 PDAC and 14 healthy control samples (Fig. 1b). This analysis revealed high expression of *TNF* (TNFα) and its receptor *TNFRSF1A* (TNFR), as well as *TNFSF12A* (TWEAK) and its receptor *TNFRSF12* (TWEAKR) (Fig. 1c, d). Other NF-κB-associated ligands and receptors, such as *TNFSF13B* (BAFF), *CD40*, and *TNFSF11* (RANKL), were also expressed, but at comparatively lower levels (Fig. S1a).

Notably, *TNF* and TWEAK (*TNFSF12*) were more highly expressed in PDAC samples compared to healthy controls (Fig. S1b, c). Furthermore, intercellular communication analysis using CellChat revealed a greater number of inferred interactions in PDAC samples relative to controls (Fig. S1d-f). Signaling pathway-specific analysis suggested an upregulation of TWEAK signaling in PDAC tumors versus healthy controls (Fig. S1f). Based on these findings, we sought to further investigate TNF-TNFR and TWEAK-TWEAKR signaling in PDAC.

To further validate these observations, we analyzed RNA-seq data from patient-derived xenografts (PDXs) and PDAC cell lines (L3.6pl and AsPC-1). Normalized counts of the NF-κB pathway ligands and receptors confirmed the predominant expression of the receptors for TNF (*TNFRSF1A)* and TWEAK (*TNFRSF12)* in tumor cells, supporting their potential roles in PDAC (Fig. 1e, f). Multiplex spectral imaging of human PDAC samples corroborated these findings, demonstrating the expression of TNFα and TWEAK in cells within the tumor microenvironment (TME) (Fig. 1g). Together, these results identify TNF-TNFR and TWEAK-TWEAKR as drivers of NF-κB signaling in PDAC.

### Cellular distribution and signaling of TNF**α** and TWEAK in PDAC

We next sought to determine the similarities and differences in the cellular source(s) of TWEAK and TNFα in PDAC. Our analysis of the scRNA-seq data revealed that immune cells, mainly macrophages, B cells, and T cells displayed *TNF* expression. In contrast, while TWEAK was also expressed in a subset of macrophages, it displayed a broader and distinct cellular expression pattern, extending to fibroblasts, endothelial, and stellate cells (Fig. 1d). In general, we observed a higher number of TWEAK expressing-cells compared to TNFα-expressing cells, suggesting it may potentially have a broader effect in PDAC tumors. To further examine and confirm these findings in vivo, we performed multiplex immunofluorescence staining for TNFα, TWEAK, CD31 (endothelial cells), α-SMA (fibroblasts), and CD68 (macrophages). Consistent with our scRNA-seq analyses, we observed TNFα staining primarily in macrophages, while TWEAK was expressed in fibroblasts and endothelial cells, as well as in some macrophages (Fig. 2a, S2a).

**Figure 2:**
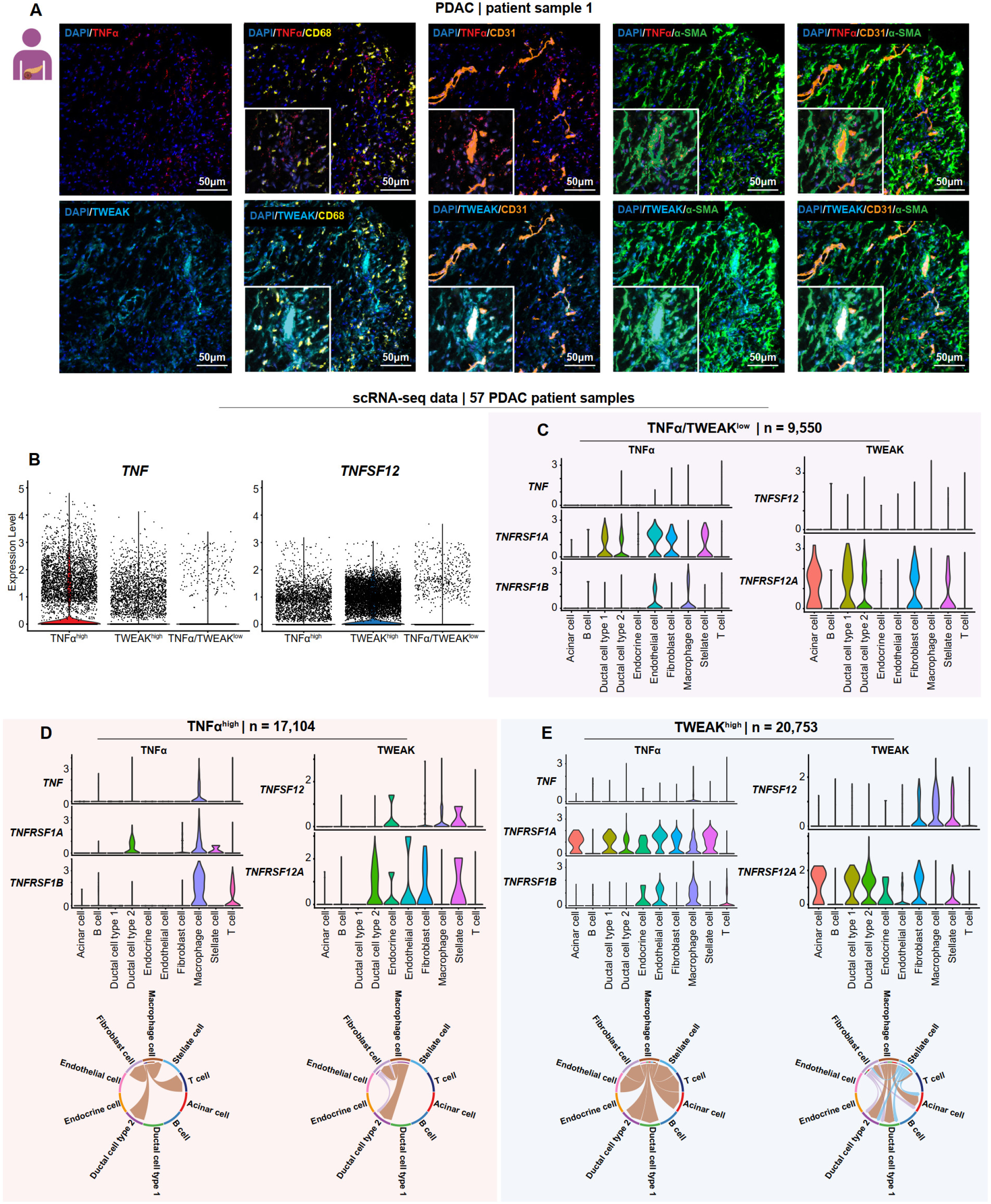
Cellular distribution and signaling networks of TNFα and TWEAK in PDAC. **A**, Multiplex immunofluorescence staining of PDAC tumor sample showing TNFα expression enriched in CD68+ macrophage-rich areas (yellow), while TWEAK expression (cyan) overlaps with CD31+ endothelial cells (orange) and α-SMA+ fibroblasts (green), indicating their distinct microenvironmental localization. Scale bar represents 50 μm (inserts: 20 μm). **B**, Expression levels of *TNF* (left) and *TNFSF12* right in PDAC patient samples grouped by TNFα/TWEAK^low^ (*n = 7*), TNFα^high^ (*n = 7* samples), and TWEAK^high^ (*n = 7*) status. Violin plots show the distribution of normalized expression levels. Each dot represents an individual cell. *TNF* expression is highest in the TNFα^high^ group, whereas *TNFSF12* expression is notably higher in the TWEAK^high^ group, reflecting a broader distribution of *TNFSF12*-expressing cells in this population. **C-E**, Expression patterns of TNFα and TWEAK ligand-receptor interactions across different cell types, analyzed in TNFα/TWEAK^low^, TNFα^high^, and TWEAK^high^ groups. CellChat analysis was used to compare TNFα and TWEAK signaling strengths, highlighting their differential roles in tumor communication networks.

To gain biological insight into the effects of TNFα and TWEAK signaling in tumor cells, we stratified PDAC samples into three groups: (1) TNFα/TWEAK^low^ (low expression of both ligands), (2) TNFα^high^(high TNFα, low TWEAK), and (3) TWEAK^high^ (high TWEAK, low TNFα) (Fig. 2b, S2b–d). CellChat analysis revealed that each group exhibited distinct ligand–receptor signaling patterns (Fig. 2c–e).

We confirmed that the TNFα/TWEAK^low^ group displayed low expression of both cytokines, and as expected, had minimal network connectivity (Fig. 2c). In the TNFα^high^ group, *TNF*–*TNFRSF1A* and *TNF*–*TNFRSF1B* interactions were predominant, with macrophages serving as the primary source of *TNF* (Fig. 2d). *TNF*–*TNFRSF1A* signaling was directed from macrophages to ductal cell type 2 and fibroblasts, whereas *TNF*–*TNFRSF1B* signaling primarily involved macrophage–T cell interactions (Fig. S2e).

In contrast, the TWEAK^high^ group displayed a broader and more complex signaling network. *TNFSF12* (TWEAK) was highly expressed in macrophages, fibroblasts, and stellate cells. Its receptor, *TNFRSF12A* (Fn14; TWEAKR), was most prominently expressed in ductal cell type 2 and fibroblasts, and was present to a lesser extent in macrophages (Fig. 2e). This pattern suggests a primarily paracrine mode of signaling, in which TWEAK produced by macrophages acts on neighboring cells with higher receptor expression. The TWEAK^high^ group exhibited a higher number of inferred interactions and stronger overall signaling activity compared to the TNF ^high^ group, engaging macrophages, fibroblasts, stellate cells, acinar cells, and ductal cells (Fig. S2f).

Collectively, these findings indicate that TWEAK signaling engages a broader range of stromal and epithelial components, likely through paracrine mechanisms, suggesting a more expansive influence of the TME. In contrast, TNFα signaling remains more restricted and immune-centric, with interactions primarily among immune and stromal cells.

### TWEAK induces a limited transcriptional program compared to TNF**α**

We next sought to characterize the functional signatures associated with TNFα^high^ and TWEAK^high^ cells by analyzing differentially expressed pathways. This revealed an upregulation of pathways involved in translational activity in TNFα^high^tumor cells and mitochondrial-related processes in tumor cells from TWEAK^high^ tumors (Fig. S3a). Collectively, these findings suggest that tumor cells from TNFα^high^ and TWEAK^high^ tumors display activation of distinct programs in response to signals from the tumor microenvironment.

To further delineate the similarities and differences between canonical and noncanonical NF-κB signaling, we performed transcriptional profiling on the PDAC cell lines AsPC-1 and L3.6pl. First, we assessed the activation dynamics of canonical vs. noncanonical NF-κB signaling by treating PDAC cell lines with TNFα or TWEAK and monitoring RELA, NFκB1 (p105/p50), NFκB2 (p100/p52) and RELB activation. As expected, RELA phosphorylation was detected within 5 min of TNFα treatment, indicating rapid activation of the canonical pathway. In contrast, processing of p105 to p50 was not observed until 8 h post-treatment. This temporal separation suggests that early canonical signaling relies on pre-existing p50, while p105 processing may serve as a delayed, secondary mechanism to sustain or reinforce NF-κB activity in response to TNFα treatment. Like NFκB1, TWEAK-induced p100 processing to p52 and RELB induction also occurred at a later time point (8 h) (Fig. 3a). Together, these findings suggest that while TNFα rapidly activates canonical NF-κB signaling through early RELA phosphorylation, TNFα-induced p105 processing and TWEAK-induced p100 processing and RELB induction occur later. This temporal pattern implies that RELA is rapidly induced by TNFα while p52/RELB activation by TWEAK is substantially delayed^10,19^.

**Figure 3:**
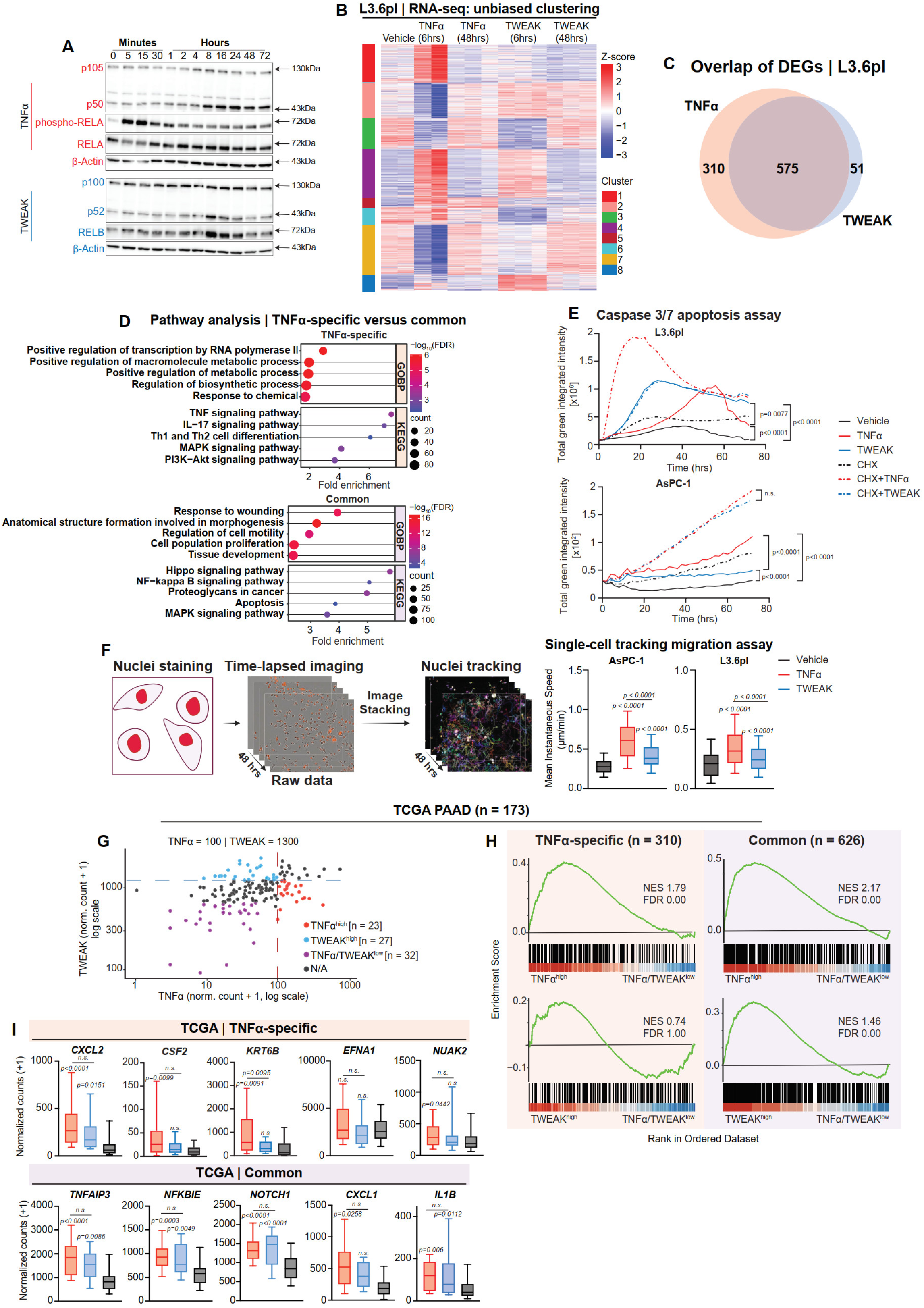
The functional differences and similarities between TNFα and TWEAK signaling activation in PDAC. **A**, Western blot analysis of p105/p50, p100/p52, phospho-RELA, and RELB following TNFα and TWEAK treatments across multiple time points (0–30 minutes and 1– 72 hours). Representative of *n = 3* independent experiments. **B**, Heatmap of differentially expressed genes following RNA-seq analysis on L3.6pl cells treated with TNFα and TWEAK for 6 and 48 hours. Unbiased clustering analysis was performed after differential expression with DESeq2. **C**, Venn diagram displaying the number of TNFα-specific, TWEAK-specific, and commonly differentially expressed genes (DEGs), derived from DESeq2 analysis and unbiased clustering analysis. Selections were based on Log2FC ≥ 1, FDR ≤ 0.05, and base mean ≥ 10. **D**, Pathway analysis for biological processes (GO) and KEGG of TNFα-specific (top) and common genes (bottom) identified in **C**. The top pathways were selected based on FDR values. **E**, Caspase 3/7 activity in L3.6pl and AsPC-1 cells over 72 hours following treatment with TNFα (10 ng/ml), TWEAK (10 ng/ml), and cycloheximide (CHX; 10 µM) as indicated. Values represent total green integrated intensity (normalized to vehicle control and cell number at each time point). Area under the curve (AUC) was calculated for each condition, and p-values were determined using unpaired t-tests based on AUC values. *n.s.* = not significant. **F**, Single-cell migration tracking assay captured every 15 minutes over 48 hours in AsPC-1 and L3.6pl cells using IncuCyte live-cell imaging (10× magnification). Boxplots depict the mean instantaneous speed calculated from migrated tracks for AsPC-1 and L3.6pl cells following vehicle, TNFα, and TWEAK treatments (p-value: one-way ANOVA; Dunnett’s multiple comparisons relative to vehicle control). **G**, Scatter plot of normalized expression levels of TNFα and TWEAK (log scale) in 173 PDAC patient samples from TCGA. Each dot represents an individual sample. Based on TNFα and TWEAK expression thresholds (TNFα ≥100 and TWEAK<1300), samples were classified as TNFα^high^ (red), TWEAK^high^ (blue; TNFα<100 and TWEAK≥1300), TNFα/TWEAK^low^ (purple; TNFα<60 and TWEAK<700, or not assigned (black). **H**, Gene Set Enrichment Analysis (GSEA) results for TNFα-specific (*n=310*) and common *(n=626*) gene sets in the TNFα^high^, TWEAK^high^, and TNFα/TWEAK^low^ expression from TCGA data. Normalized Enrichment Scores (NES) and False Discovery Rate (FDR) values are shown in each plot. TNFα^high^ samples show significant enrichment of both TNFα-specific and common gene sets, whereas TWEAK^high^ samples only display enrichment of the TNFα-specific gene set. **I**, Boxplots showing expression of selected genes from TNFα-specific and common gene sets in PDAC samples grouped by TNFα^high^, TWEAK^high^, and TNFα/TWEAK^low^ status. Genes were selected based on their rankings from the GSEA analysis. TNFα-specific genes (top panels) show higher expression in TNFα^high^ samples, while common genes (bottom panels) are elevated in both TNF ^high^ and TWEAK^high^ samples. p-values are shown for significant differences (unpaired t-tests).

To capture early and late response genes, we performed RNA-seq at 6- and 48-h post-treatment in AsPC-1 and L3.6pl cells (Fig. 3b, S3b). Unbiased clustering analysis identified eight gene clusters, where TNFα-specific clusters (1, 5) were enriched in inflammation, immune response, metabolic reprogramming, cell migration, and ECM organization (Fig. S3c, S3d), while shared TNFα/TWEAK clusters (4) were associated with cellular homeostasis, growth, and division (Fig. S3d). Notably, the small TWEAK-dominant cluster (8) exhibited a similar enrichment with TNFα-specific cluster 1 and 5 (Fig. S3d, e), indicating that TWEAK does not drive a distinct transcriptional program in PDAC cells but rather functions to regulate a subset of genes also controlled by TNFα. Additional clusters were linked to cell cycle (cluster 2; Fig. S3f), developmental reprogramming (cluster 3; Fig. S3g), structural organization and metabolism (cluster 6; Fig. S3h), and DNA repair and proliferation (cluster 7; Fig. S3i).

To further confirm the role of TWEAK in inducing a subset of TNFα-regulated genes, we performed differential expression analysis and identified a number of TNFα-specific (n = 310) and commonly regulated (n = 575) genes, but only a small subset of genes preferentially activated by TWEAK (n = 51) (Fig. 3c). These findings demonstrate that TWEAK signaling does not elicit a distinct transcriptional response in PDAC cells but rather modulates core cellular homeostasis and growth pathways also activated by TNFα.

### TNF**α** and TWEAK both promote apoptosis and migration in PDAC

To further delineate the similarities and differences between TNFα and TWEAK, we analyzed GO enrichment of their shared gene set (Fig. 3d). The overlap indicates that both cytokines regulate genes associated with proliferation, migration, apoptosis, and tissue remodeling. To functionally validate these observations, we examined apoptosis and migration in response to TNFα and TWEAK treatment. Caspase-3/7 activity assays confirmed that the apoptotic machinery was functional in both cell lines following treatment with the positive control staurosporine (Fig. S4a)^20^. TNFα and TWEAK both significantly induced low levels of apoptosis in L3.6pl cells, albeit with different time kinetics, but did not appreciably affect apoptosis in AsPC-1 cells (Fig. 3e). As previously described^21^, inhibition of protein synthesis with cycloheximide exacerbates the pro-apoptotic effect of TNF receptor family activation. Consistently, concomitant cycloheximide treatment invoked similar pro-apoptotic effects of TNFα and TWEAK in AsPC-1 cells and facilitated a more rapid and pronounced induction of apoptosis by TNFα in L3.6pl cells, while TWEAK similarly induced apoptosis independent of inhibition of protein synthesis in L3.6pl cells. Collectively, these findings highlight that TNFα and TWEAK are both capable of inducing apoptosis in PDAC cells in a protein synthesis-regulated manner.

In our previous work we identified an important role for TNFα-induced NF-κB signaling in promoting PDAC cell migration^4^. Thus, we performed single-cell migration assays to compare the effects of TNFα and TWEAK treatment on cell migration. Consistent with our previous work, TNFα significantly enhanced migration in both cell lines, while TWEAK had a significant, but more modest effect (Fig. 3f, S4a).

### TNF**α**- and TWEAK-associated gene signatures in patient PDAC tissues

To confirm the TNFα-specific and common gene sets we identified from our transcriptional analysis and to assess clinical relevance, we utilized bulk RNA-seq data from TCGA pancreatic cancer samples stratified into TNFα^high^, TWEAK^high^, and TNFα/TWEAK^low^ groups (Fig. 3g). Differential expression analysis revealed that TWEAK^high^ samples exhibited a markedly higher number of upregulated genes (compared to TNFα/TWEAK^low^) than TNFα^high^ samples (1063 vs. 57, respectively), suggesting that TWEAK drives a broader transcriptional response in patient PDAC tumors (Fig. S4b). Consistent with our in vitro studies, we observed significant enrichment of the common gene set in both TNFα^high^ and TWEAK^high^ groups, whereas only TNF ^high^ samples showed significant enrichment of the TNFα-specific gene set (Fig. 3h). Selected genes from these sets further validated these observations, with TNFα-specific genes expressed predominantly in TNF ^high^ samples and common genes enriched in α and TWEAK groups (Fig. 3i). Together, these results validate the TNFα-specific and common gene sets we identified and demonstrate their relevance in human PDAC tissues.

### TNF**α** and TWEAK differentially activate NF-**κ**B binding dynamics

Although NF-κB signaling is a well-established mediator of gene activation, the specific roles of the canonical (TNFα) and noncanonical (TWEAK) pathways remain underexplored. To investigate this, we analyzed the genome-wide binding dynamics of RELA and RELB, the respective downstream transcription factors activated by TNFα and TWEAK respectively (Fig. 1a). scRNA-seq analysis confirmed RELA and RELB expression in PDAC cells (Fig. 4a). Notably, multiplex immunofluorescence staining in PDAC patient samples confirmed that activation of RELA (via phosphorylation) and expression of RELB in epithelial tumor cells (expressing cytokeratin) is closely associated with the expression of the respective ligand in the proximal TME in patient tumor samples (Fig. 4b).

**Figure 4:**
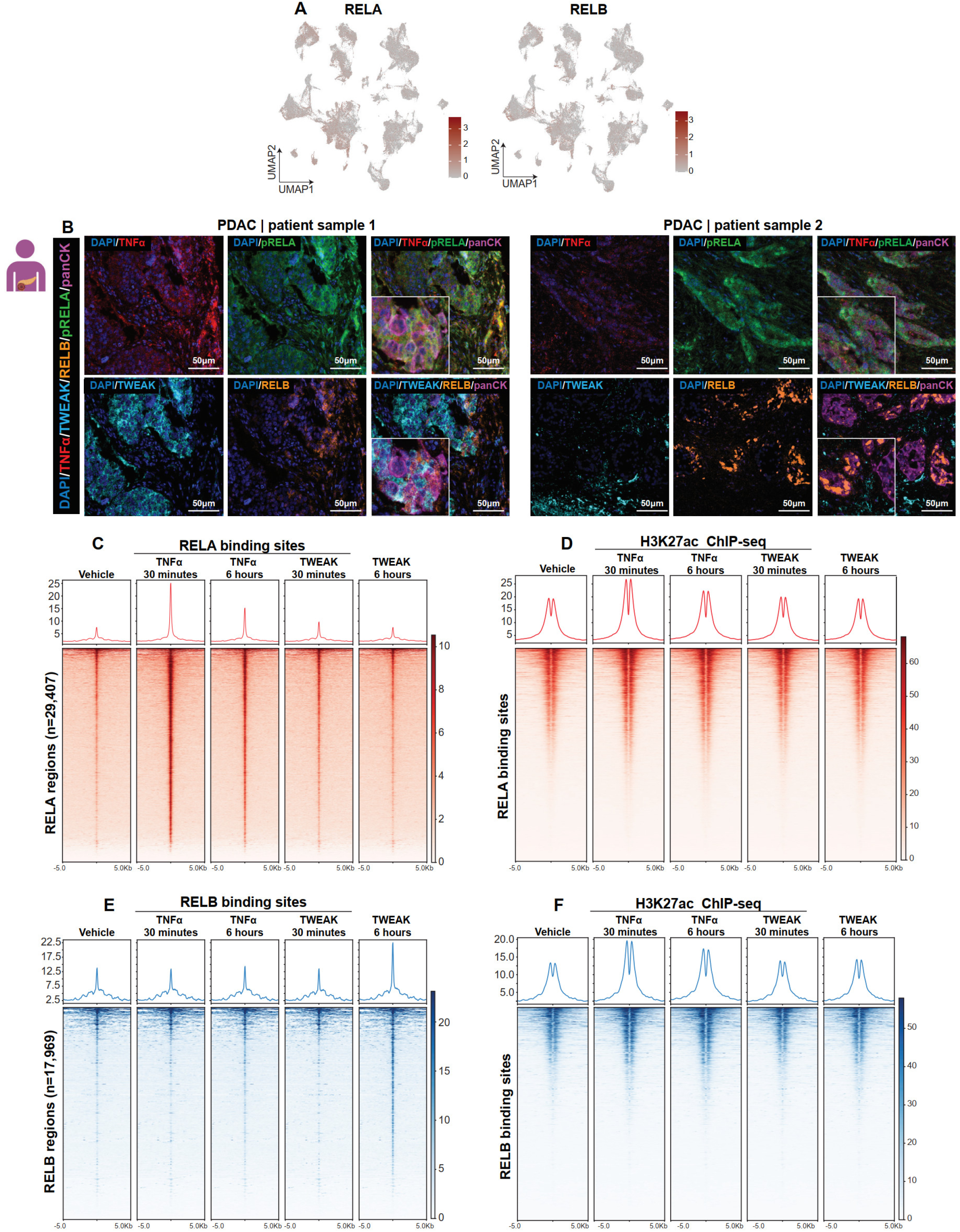
Epigenetic mechanism of canonical and noncanonical NF-κB signaling in PDAC. **A**, UMAP plots showing the expression patterns of phospho-RELA and RELB in PDAC patient samples (*n = 136,163 cells, n = 57 donors*) **B**, Multiplex immunofluorescence staining of PDAC patient samples showing RELA (green) and RELB (orange) expression in TNFα (red) and TWEAK (cyan) expressing cells. **C&D**, Heatmaps and signal intensity profiles of ChIP-seq data for RELA (**C**) and H3K27ac (**D**) across identified RELA binding sites (*n=29,407*) in PDAC cells. Vehicle-treated and TNFα- or TWEAK-stimulated samples (30 minutes and 6 hours) are shown. RELA ChIP-seq reveals rapid binding of RELA after TNFα treatment (30 minutes), while TWEAK treatment shows no RELA recruitment. H3K27ac ChIP-seq demonstrates that TNFα treatment induces strong H3K27ac occupancy at RELA sites, with a more robust effect at 30 minutes compared to 6 hours. **E&F**, Heatmaps and signal intensity profiles of ChIP-seq data for RELB (**E**) and H3K27ac (**F**) across identified RELB binding sites (*n=17,969*) in PDAC cells treated with vehicle, TNFα, or TWEAK for 30 minutes and 6 hours. RELB ChIP-seq shows delayed recruitment by TWEAK (6 hours) but no significant binding after TNFα stimulation. H3K27ac ChIP-shows minimal changes in H3K27ac occupancy at RELB sites.

To investigate the temporal recruitment of RELA and RELB to chromatin and associated epigenetic changes, we performed ChIP-seq for RELA, RELB, and H3K27ac following TNFα or TWEAK treatment for 30 min and 6 h, based on activation kinetics observed in Fig. 3a. TNFα treatment induced selective RELA (but not RELB) binding at early time points, accompanied by an increase in adjacent H3K27ac occupancy (Fig. 4c, d), while TWEAK treatment selectively induced RELB (but not RELA) binding at a later time point without notably increasing H3K27ac at these sites (Fig. 4e, f). These findings demonstrate that TNFα specifically activates RELA and promotes H3K27ac enrichment at NF-κB-associated regions, whereas TWEAK selectively activates RELB without significantly altering H3K27ac occupancy.

### TNF**α** induces broader NF-**κ**B binding and epigenetic activation than TWEAK

To investigate transcriptional regulatory mechanisms controlled by canonical and noncanoical NF-κB signaling, we performed differential binding analysis (DiffBind) comparing RELA and RELB binding at 30 min following TNFα treatment and 6 h after TWEAK treatment, respectively. This analysis revealed that RELA bound significantly more regions than RELB (Fig. 5a, S5a). Consistent with our RNA-seq results where TNFα induced a distinct subset of genes, while TWEAK-induced a subset of genes that was also induced by TNFα (Fig. 3c), RELB-bound regions were also bound by RELA, while many RELA-bound regions displayed no significant enrichment of RELB (Fig. 5a-b). Thus, we could separate RELA/RELB-bound regions into cluster 1 containing RELA-specific regions and cluster 2 containing commonly bound regions (Fig. 5b-d).

**Figure 5:**
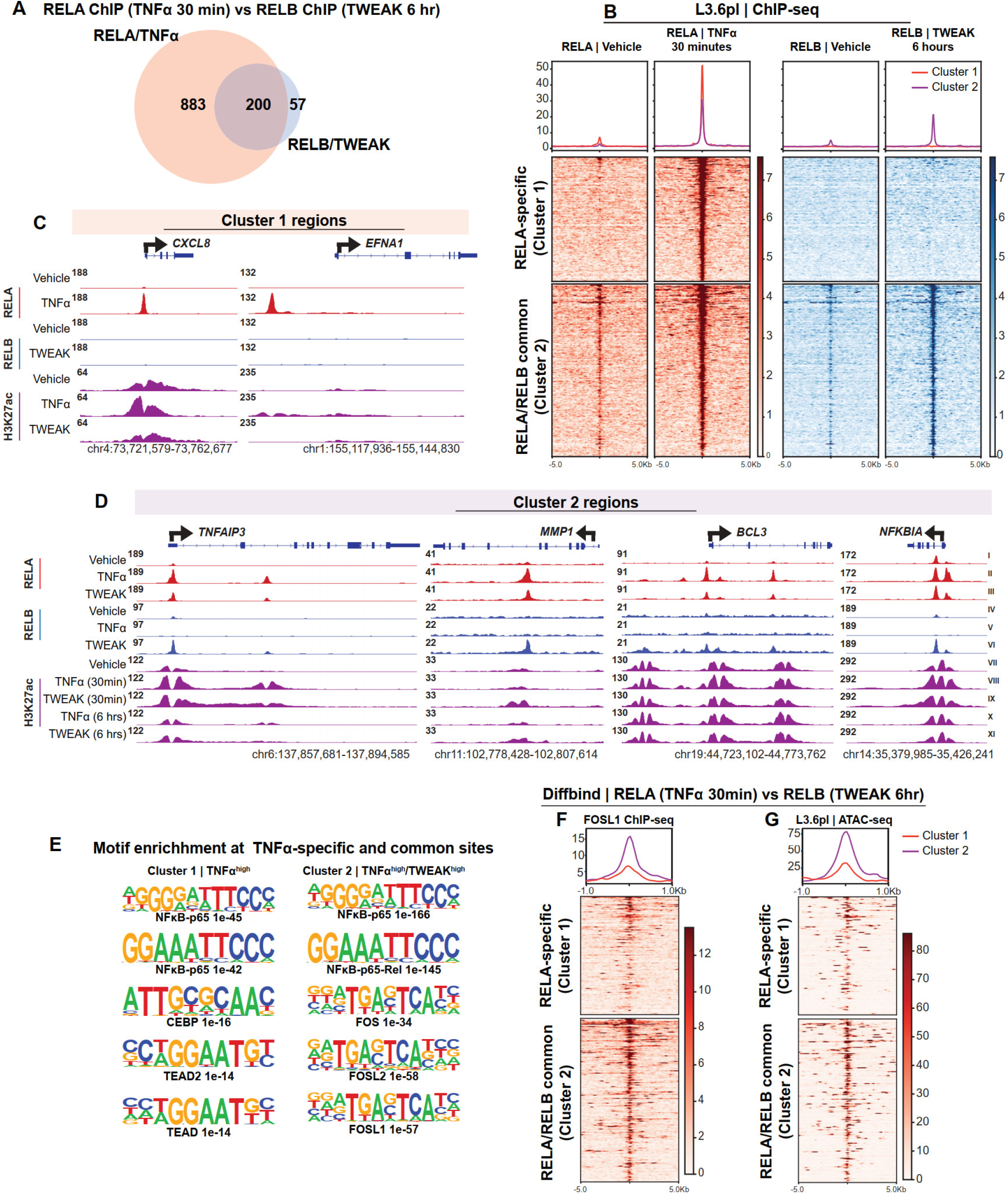
RELA/RELB common regions are associated with chromatin accessibility and AP-1 factors. **A**, Venn diagram comparing RELA-specific and RELB-specific binding regions identified by DiffBind and bedtools intersect analysis. RELA binding was assessed 30 minutes after TNFα treatment, while RELB binding was assessed 6 hours after TWEAK treatment. The regions were selected based on: RELA/TNFα-specific regions (Log2FC ≥ 1, FDR ≤ 0.05). RELB/TWEAK-specific regions (FDR ≤ 0.05, with only 57 regions differentially bound). Common regions (FDR ≤ 0.9 and ≥ -0.9). **B**, Heatmap of RELA-specific and common regions identified in **D**, along with corresponding signal comparisons across treatments. RELA and RELB signals for the 57 TWEAK-specific regions were included in the common regions (*n = 257*) following vehicle, TNFα, and TWEAK treatments, reflecting shared regulatory elements between canonical and noncanonical NF-κB signaling. **C-D**, Genome browser (IGV) tracks of ChIP-seq data in L3.6pl cells showing RELA, RELB, and H3K27ac signals at: RELA/TNFα-specific gene loci (**C**; *CXCL8* and *EFNA1*) and Common gene loci (**D**; *TNFAIP3*, *MMP1*, *BCL3*, and *NFKBIA*). Tracks illustrate RELA, RELB, H3K27ac signals following vehicle, TNFα, and TWEAK treatments, highlighting distinct and overlapping regulatory regions of RELA and RELB. **E**, Top enriched motifs identified in Cluster 1 (RELA-specific regions) and Cluster 2 (RELA/RELB common regions) using HOMER motif analysis, demonstrating distinct transcription factor binding preferences. **F**, Heatmap of FOSL1 signals across Cluster 1 (RELA-specific) and Cluster 2 (RELA/RELB common regions), showing its preferential binding patterns. **G**, ATAC-seq signal intensity at Cluster 1 and Cluster 2 in L3.6pl cells, indicating differences in chromatin accessibility between RELA-specific and RELA/RELB common regions.

Analysis of H3K27ac occupancy at these regions confirmed the more pronounced effect of TNFα-activated RELA in promoting H3K27 acetylation at NF-κB-associated loci (Fig. S5b). These findings demonstrate that TWEAK-induced noncanonical NF-κB selectively engages RELB with limited epigenetic activation at a subset of regions displaying RELA binding after activation of canonical NF-κB signaling by TNFα treatment. Additionally, TNFα stimulated RELA binding to a number of additional regions selectively bound by RELA, but not RELB, which also display enhanced H3K27ac occupancy following ligand treatment. This differential regulation underscores the distinct yet overlapping contributions of TNFα and TWEAK in shaping NF-κB-dependent epigenetic activation of transcription.

### RELB relies on chromatin accessibility and AP-1 factors for gene regulation

To further explore the regulatory mechanisms associated with the differential transcriptional effects elicited by RELA and RELB downstream of TNFα and TWEAK signaling, respectively, we examined transcription factor enrichment on the RELA-specific genomic regions (cluster 1) as well as the RELA/RELB-common regions (cluster 2) (Fig. 5b). Cluster 1 was primarily enriched for RELA and NFκB1, while in addition to enrichment for REL factors, cluster 2 also displayed a significant enrichment of AP-1 factors (e.g., FOS and JUN families of AP-1) (Fig. S5c). Similarly, motif enrichment analysis confirmed the presence of RELA (NFκB-p65) and TEAD motifs in cluster 1, whereas cluster 2 displayed a significant enrichment of AP-1 motifs in addition to RELA motifs (Fig. 5e). These results suggest that TNFα-induced RELA is able to largely function independently, while TWEAK-induced RELB may require additional transcription factors, such as AP-1 factors, for activity.

To further test this, we examined FOSL1 occupancy on cluster 1 (RELA-specific) and cluster 2 (RELA/RELB common regions). Intersection analysis revealed that 77% of RELB regions overlapped with FOSL1 (Fig. S5d) while just 27% of RELA-occupied regions intersected with FOSL1. Our previous published data^4^ indicated that RELA-FOSL1 co-bound regions regulate cell migration in PDAC through the convergence of MAPK and NF-κB signaling pathways. Consistently, FOSL1 binding was higher in common regions (Cluster 2) than in RELA-specific regions (Cluster 1) (Fig. 5f, S5e).

We next sought to test whether the differences in RELA and RELB binding may be related to differences in chromatin accessibility prior to pathway activation. Thus, we examined chromatin accessibility by ATAC-seq^22^ and observed that common regions (Cluster 2) exhibited significantly higher chromatin accessibility compared to RELA-specific regions (Cluster 1) (Fig. 5g, S5e). These findings underscore that noncanonical TWEAK signaling through RELB relies heavily on an open chromatin state to regulate gene expression, while TNFα-driven RELA binding can occur even at closed chromatin regions. Together our results reveal that TNFα-driven RELA autonomously binds and regulates gene expression by increasing H3K27ac and activating enhancer regions, while TWEAK-induced RELB relies on the binding of other factors such as AP-1 to provide an open chromatin conformation and active epigenetic state. Thus, these two pathways function with partially overlapping, but distinct mechanisms that may be useful for dissecting their individual contributions to cancer biology and potentially targeting individual aspects of NF-κB biology.

## Discussion

NF-κB signaling is a key driver of PDAC, with persistent activation observed in 70% of tumors^1,23^. While the canonical (TNFα-TNFR) NF-κB pathway has been extensively studied, significantly less is known about the noncanonical (TWEAK-TWEAKR) pathway or the functional and differential epigenetic mechanisms by which these two pathways function in PDAC. Through multi-omics approaches, we uncover divergent chromatin-binding dynamics of RELA and RELB, highlighting the ability of TNFα to remodel chromatin and autonomously activate NF-κB-dependent transcription through RELA, while TWEAK-driven RELB activation requires pre-existing chromatin accessibility and co-factors such as AP-1.

Our results demonstrate that TNFα rapidly activates canonical NF-κB signaling, with RELA driving the expression of a vast number of genes. Functionally, TNFα-induced apoptosis is significantly enhanced when translation is inhibited, suggesting a dependency on rapidly synthesized survival factors such as MCL-1 and XIAP^20,24,25^. In contrast to the rapid transcriptional effects of TNFα, TWEAK-induced noncanonical NF-κB signaling exhibits slower activation dynamics and lacks a unique transcriptional program in PDAC cells, with nearly all induced genes also being induced by TNFα.

Our scRNA-seq analysis revealed that TWEAK is expressed by a broader range of cell types in PDAC whereas TNFα is predominantly expressed by immune cells. Consistent with this, CellChat-based interaction analysis showed that TWEAK has greater cumulative signaling strength and a more complex network of ligand–receptor interactions. Consistent with a broader expression of TWEAK in the TME, TCGA analysis identified more drastic transcriptional differences in TWEAK^high^ tumors compared to TNFα^high^tumors, suggesting that the TWEAK signal produced by more cells in the TME is able to engage more individual tumor cells and elicit an overall stronger transcriptional response in bulk RNA-sequencing. This finding is consistent with our multiplex immunofluorescence stainings showing a broad expression of TWEAK in the PDAC TME. Interestingly, in contrast to the in vivo data, TNFα stimulation of PDAC cell lines in vitro induced a more robust transcriptional response than TWEAK. Thus, these findings support a model in which TWEAK recapitulates many of the transcriptional effects of TNFα, but through a slower, broader, and more distributed signaling program within the TME. This is likely driven by TWEAK’s expression in diverse stromal and epithelial compartments. In contrast, TNFα signaling elicits a significantly stronger effect but is more tightly controlled through interactions with specific components of the TME.

NF-κB is known to shape the epigenetic landscape in cancer by influencing chromatin accessibility, histone modifications, and transcription factor recruitment^1,5,26^. Our ChIP- and ATAC-seq analyses reveal key differences in RELA and RELB chromatin binding and activation dynamics. TNFα-induced RELA is able to bind to both open and closed chromatin regions and to establish active enhancers but promoting H3K27ac. This suggests that TNFα-driven RELA activation is sufficient to remodel chromatin and initiate transcription, reinforcing its autonomous role in NF-κB-driven oncogenic programs. In contrast, TWEAK-induced RELB binding is highly dependent on pre-existing chromatin accessibility. Unlike RELA, RELB does not promote H3K27ac, but rather binds to regions already bearing the active mark. Thus, in contrast to TNFα-driven RELA activity, TWEAK-driven RELB activation lacks autonomous chromatin-remodeling capacity. Instead, RELB requires pre-existing accessible chromatin and co-factors, such as AP-1 (FOSL, JUN) for transcriptional activity.

The identification of TNFα and TWEAK as dominant NF-κB signaling axes in PDAC presents potential therapeutic opportunities. Given the ability of TNFα to activate genes controlled by both canonical and noncanonical NF-κB pathways, strategies specifically targeting transcriptional mechanisms controlling these genes (e.g., chromatin remodeling) could potentially disrupt PDAC progression. Furthermore, the reliance of TWEAK-induced RELB on chromatin accessibility, AP-1, and possibly other co-factors suggests potential vulnerabilities that could be exploited using inhibitors of chromatin remodeling or transcription factor modulators. AP-1 has been implicated in pancreatic cancer invasion and metabolic reprogramming, making it a potential target for combination therapies with NF-κB inhibitors^27,28^. Targeting the NF-κB/AP-1 axis, either through chromatin remodeling inhibitors or MAPK blockade, could disrupt tumor adaptation mechanisms.

Our study provides a comprehensive analysis of TNFα- and TWEAK-mediated NF-κB signaling in PDAC, revealing distinct transcriptional, epigenetic, and functional differences. Our data suggests that TNFα is a primary driver of inflammation, metabolic adaptation, and migration through direct chromatin remodeling and autonomous RELA activity, whereas TWEAK relies on chromatin accessibility and transcriptional co-factors such as AP-1 to regulate gene expression. These findings suggest that TNFα-driven functions in PDAC may be differentially vulnerable compared to TWEAK-driven effects, which may be targeted by disrupting NF-κB/AP-1 interactions. Moreover, these results provide important new insights into the similarities and differences between canonical and noncanonical NF-κB and reveal important differences in the effects elicited by different components of the TME on PDAC tumor cell gene expression. Further studies will be necessary to elucidate how these mechanisms can be exploited for improved patient therapy.

## Materials and methods

All details are in supplementary materials and methods.

### Statistics and Reproducibility

All images presented are representative of at least three biological replicates, except for single-cell migration assays, where graphs represent approximately 3,000 cells. Statistical significance was determined using unpaired t-tests. Statistical analysis of apoptosis curves was performed by calculating the area under the curve (AUC) followed by unpaired t-tests of the AUC values. Box plots display the 10th to 90th percentile range, with error bars representing the spread of the central 80% of values; these are percentile-based and do not correspond to standard deviation (s.d.), standard error of the mean (s.e.m.), or confidence intervals (c.i.). Upregulated genes and regions were identified using thresholds of log2 fold change >1, false discovery rate (FDR) ≤0.05, and BaseMean >10. All graphs and statistical analyses were conducted using GraphPad Prism 10.4.1.

### Schematic images

Model figures were generated using https://www.biorender.com/.

## Data Availability Statement

All raw and processed data are available under the GEO accession numbers GSE296831 and GSE296832.

## Code Availability

Custom software was used to perform automated cell tracking and trajectory analysis and is freely available at: https://github.com/Clark-Lab-Stuttgart/incucyte_cell_tracking.

## Compliance with ethics statement

This study received all necessary ethical approvals from the Institutional Review Board (IRB) under protocols [354-06, 66-06, and 19-012104]. All experimental procedures and methods were conducted in accordance with relevant institutional guidelines and regulations. Informed consent was obtained from all human participants or their legal guardians prior to inclusion in the study, in accordance with IRB-approved protocols and institutional standards.

## Conflict of interest

There are no conflicts of interest.

## Funding

Research was supported by the Robert Bosch Stiftung.

## Notes

https://www.ncbi.nlm.nih.gov/geo/query/acc.cgi?acc=GSE296831

https://www.ncbi.nlm.nih.gov/geo/query/acc.cgi?acc=GSE296832

### Competing Interest Statement

The authors have declared no competing interest.

