## Supplementary Materials and Methods for "Epigenetic Context Defines the Transcriptional Activity of Canonical and Noncanonical NF-κB Signaling in Pancreatic Cancer"

Aggrey-Fynn et al.

Supplementary Figure Legends

Supplementary Materials and Methods

Supplementary References

Supplementary material 1: TNFα-specific and common genes

Supplementary material 2: Western blots used in the study (L3.6pl)

**Supplementary figure legends**

**Supplementary figure 1: NF-κB signaling is elevated in PDAC tumor cells compared to normal cells. A**, UMAP plots displaying the expression patterns of BAFF (*TNFSF13B*), CD40, and RANKL (*TNFSF11*) ligands, highlighting their distribution across cell populations in PDAC patient samples. **B & C**, UMAP plots and violin plots comparing the expression levels of *TNF* (TNFα) and *TNFSF12* (TWEAK) between PDAC and healthy patient samples, demonstrating increased NF-κB ligand expression in tumor cells. **D**, Circle plots illustrating intercellular communication networks between cells from healthy and PDAC patient samples. Blue lines indicate reduced communication in PDAC, while red lines indicate increased communication relative to healthy cells. Arrows denote the directionality of intercellular signaling interactions. **E**, Comparison of NF-κB pathway signaling flow between PDAC and healthy cells, calculated as the summation of ligand-receptor gene expression products across all sender-receiver pairs. Blue indicates increased communication in PDAC, while red indicates decreased communication relative to healthy samples. **F**, Bar graph quantifying total inferred interactions and interaction strength between healthy and PDAC samples, further emphasizing the enhanced NF-κB signaling activity in tumor cells.

**Supplementary figure 2: Cellular distribution and signaling networks of TNF and TWEAK in PDAC. A**, Multiplex immunofluorescence staining of PDAC tumor sample 2 showing TNFα expression enriched in CD68+ macrophage-rich areas (yellow), while TWEAK expression (cyan) overlaps with CD31+ endothelial cells (orange) and α-SMA+ fibroblasts (green). Scale bar represents 50 μm (inserts: 20 μm). **B**, Violin plots display the expression levels of TNFα and TWEAK across all PDAC patient samples, ranked from highest to lowest expression. **C**, Dot plot showing **TNF** and **TNFSF12** expression across PDAC patient samples grouped by TNFα^high^ (red), TWEAK^high^ (blue), and TNFα/TWEAK^low^ (purple) status. Each dot represents the percent of cells expressing the gene within a sample (dot size) and the average expression level (dot color, log2 scale). **D**, UMAP plots showing the expression patterns of ***TNF*** and ***TNFSF12* in the** TNFα^high^, TWEAK^high^, and TNFα/TWEAK^low^ groups. **E**, The directionality of intracellular communication mediated by *TNF*-*TNFRSF1A* and *TNF*-*TNFRSF1B* ligand-receptor interactions is shown in TNFα^high^ patient samples. **F**, A bar graph compares the total inferred interactions and interaction strength between TNFα^high^ and TWEAK^high^ patient groups, emphasizing differences in signaling activity. **G-H**, Scatter plots visualize the intensity of outgoing and incoming interactions across a two-dimensional manifold, where the size of each circle represents the number of significantly expressed receptor-ligand pathways across the two sample groups.

**Supplementary figure 3: The functional differences and similarities between TNFα and TWEAK signaling activation in PDAC. A**, Pathway analysis for biological processes (GO) and KEGG of enriched genes in TNFα^high^ and TWEAK^high^ versus TNFα/TWEAK^low^ samples, highlighting distinct functional pathways activated by each signaling axis. The top pathways were selected based on FDR values. **B**, Heatmap of differentially expressed genes following RNA-seq analysis on AsPC-1 cells treated with TNFα and TWEAK for 6 and 48 hours. Unbiased clustering analysis was performed after differential expression with DESeq2, identifying distinct transcriptional responses to TNFα and TWEAK signaling. **C**, Pathway analysis for biological processes (GO) and KEGG for genes in clusters 1, 3 & 7, and 6, highlighting key functional pathways enriched in each group. The top pathways were selected based on FDR values. **D-I**, Pathway analysis for biological processes (GO) and KEGG for genes in clusters 1–8, identified through unbiased clustering analysis. The selected pathways represent the most significantly enriched biological functions and signaling mechanisms associated with each cluster, based on FDR values.

**Supplementary figure 4: The functional differences and similarities between TNFα and TWEAK signaling activation in PDAC. A**, Caspase 3/7 activity in L3.6pl and AsPC-1 cells over 72 hours following treatment with staurosporine (STS; 0.1 µM). **B**, Generated migration tracks of individual AsPC-1 and L3.6pl cells following vehicle, TNFα, and TWEAK treatments for 48 hours, captured using live-cell imaging (IncuCyte, magnification = 10×). The tracks illustrate differences in cell motility and migration behavior in response to TNFα and TWEAK signaling. **C**, Volcano plots showing differentially expressed genes (DEGs) in TCGA PDAC samples identified using limma. Plots compare TNFα^high^ versus TNFα/TWEAK^low^ (left) and TWEAK^high^ versus TNFα/TWEAK^low^ (right). Red and blue points represent significantly upregulated (log2FC > 1) and downregulated (log2FC < -1) genes, respectively (adjusted p-value < 0.05). Pathway analysis for biological processes (GO) and KEGG for differentially expressed genes (DEGs) in TNFα^high^ versus TNFα/TWEAK^low^ (top) and TWEAK^high^ versus TNFα/TWEAK^low^ (bottom) PDAC patient samples from TCGA. TWEAK^high^ samples show overlapping set of enriched pathways.

**Supplementary figure 5: RELA/RELB common regions are associated with chromatin accessibility and AP-1 factors. A**, Principal Component Analysis (PCA) comparing RELA/TNFα and RELB/TWEAK binding profiles following DiffBind analysis (top), showing distinct clustering of transcription factor occupancy. ChIP-seq data were obtained from triplicate samples. Bottom, a MA plot displays the differentially bound regions identified by DiffBind analysis, highlighting a significantly higher number of RELA/TNFα-bound regions compared to RELB/TWEAK-bound regions, suggesting broader chromatin engagement by RELA. **B**, Heatmap of H3K27ac signals in RELA-specific and common binding regions, identified in DiffBind analysis, with corresponding signal comparisons across vehicle, TNFα, and TWEAK treatments. **C**, Dot plot showing the top transcription factors enriched in RELA-specific regions (Cluster 1, right) and RELA/RELB common regions (Cluster 2, left), identified through ChIP-Atlas analysis. Transcription factors were selected based on FDR values, highlighting differential regulatory factor associations. **D**, Venn diagram illustrating the overlap between FOSL1, RELA, and RELB binding regions, generated through bedtools intersect analysis. The diagram highlights unique and shared genomic loci, reinforcing the role of AP-1 factors in NF-κB signaling integration. **E**, IGV tracks displaying RELA, RELB, FOSL1, ATAC-seq, and H3K27ac signals (L3.6pl) at clusters 1 & 2, providing insight into transcription factor binding, chromatin accessibility, and histone acetylation dynamics.

**Materials and Methods**

**Supplementary Materials and Methods**

**Cell Culture**

AsPC-1 (RRID:CVCL_0152) cells were maintained in RPMI 1640 Medium (Corning). L3.6pl (RRID:CVCL_0384) cells were maintained in phenol red-free minimum essential media (MEM; Thermo Fischer Scientific). Media were supplemented with 10% FBS (Corning), 1% Penicillin/streptomycin (Thermo Fischer Scientific), and 1% L-Glutamine (Corning, for MEM media). Cells were split upon reaching 70-80% confluence. All treatments were performed in the appropriate media and the list of proteins and inhibitors and concentrations used are provided in Supplementary Table S1. Cells were treated with TNF (10ng/ml; R&D systems 210-TA-100), TWEAK (10ng/ml; R&D systems 1090-TW-025), caspase 3/7 (1:500; ThermoFisher; C10432), and cycloheximide (10 µM; Sigma Aldrich; C7698),

**Protein isolation, RNA-seq, and ChIP-seq library preparation**

Protein isolation and western blots were performed as previously reported ^1^. The following primary antibodies were used at the indicated dilutions: phospho-NFκB p65 (S536) (93H1) (phospho-RELA; 1:1000; Cell Signaling Technology; 3033; RRID:AB_331284), RELB (D7D7W) (1:1000; Cell Signaling Technology; 10544; RRID:AB_2797727), NF-κB2 p100/p52 (18D10) (1:1000; Cell Signaling Technology; 3017; RRID:AB_10697356), NF-κB1 p105/p50 (D4P4D) (1:1000; Cell Signaling Technology; 13586; RRID:AB_2665516), and β-Actin (D6A8) (1:1000; Cell Signaling Technology; 8457; RRID:AB_10950489). Secondary antibodies were used at a 1:2500 dilution and included goat anti-rabbit IgG Starbright Blue 700 (Bio-Rad; 1200416; RRID:AB_2721073) and goat anti-mouse IgG Starbright Blue 520 (Bio-Rad; 12005866; RRID:AB_2934034).

For the RNA-seq library, RNA quality was validated by gel electrophoresis. We used 500ng to make the libraries in triplicates for each condition. Libraries for cells were made using the TruSeq RNA Library Prep Kit V3 (Illumina) according to the manufacturer’s instructions. ChIPs and ChIP-seq were performed as previously described with minor changes ^2,3^. Libraries were prepared using the MicroPlex Library Preparation Kit v2 (Diagenode) according to the manufacturer’s protocol. Details protocol for RNA- and ChIP-seq are provided in Supplementary information. DNA quality of the resulting DNA was measured using the High Sensitivity DNA Kit (Agilent) on the Agilent TapeStation 4150 (RRID:SCR_019393). Antibodies used for ChIP are provided in the ChIP-seq methods section below. Samples were sequenced (paired-end 50 bp) on a NextSeq 2000 sequencer (P2, Illumina; RRID:SCR_023614) at the Robert Bosch Center for Tumor Diseases (RBCT).

**Cell migration analysis from incuCyte time-lapse imaging**

A total of 3000 cells were seeded and treated overnight with Nuclight Red dye (1:2000; Sartorius). The cells were then treated with proteins and inhibitors as specified in the cell culture section above. Live cell imaging was performed using the Sartorius IncuCyte (RRID:SCR_023147). Images were captured every 15 minutes for 48 hours and processed using the Basic Analyzer tool (Sartorius).

Preprocessing and image analysis were performed using FIJI (RRID:SCR_002285) ^4^. Raw brightfield and fluorescent nuclear tiff images from IncuCyte imaging were compiled using a custom script written in the ImageJ (RRID:SCR_003070) macro language ^5^. Imaging drift was corrected in batch using the Correct 3D Drift plugin ^6^. Nuclear segmentation and tracking were performed in batches using a custom Python (RRID:SCR_008394) script in ImageJ. Image nuclei were automatically segmented for each imaging frame using the StarDist2D ImageJ plugin with the included “Versatile (fluorescent nuclei)” model ^7,8^. The resulting labeled images were then tracked frame-by-frame in a semi-automated manner using the TrackMate plugin for ImageJ with a Simple Sparse LAP Tracker and a maximum linking distance and gap closing distance of 50 px ^9^. The resulting cell trajectories were output to an xml file and further analyzed using custom Python (v3.9) software utilizing the following packages: numpy (RRID:SCR_008633), scipy (RRID:SCR_008058), pandas (RRID:SCR_018214), and matplotlib (RRID:SCR_008624) ^10-13^. Tracks containing <3 time points were disregarded. Mean instantaneous speeds, mean squared displacements (MSDs), and directional correlation between trajectories were calculated as previously described ^14^. Data from different conditions was compiled, and custom Python scripts were used to perform statistical analysis and visualization.

Box plots were plotted using the 10th to 90th percentile range. The error bars represent the 10th and 90th percentiles of the data, reflecting the spread of the central 80% of values. These are percentile-based and do not correspond to standard deviation (s.d.), standard error of the mean (s.e.m.), or confidence intervals (c.i.).

**scRNAseq data analysis for publicly available datasets**

For the single cell RNAseq analysis, the following publicly available PDAC patient datasets were downloaded from the GEO database (RRID:SCR_005012), GSE154778 ^15^, GSE111672 ^16^, PRJCA001063 ^17^, GSE155698 ^18^, GSM4293555 from GSE141017 ^19^, and scRNA-seq data from the study by Chijimatsu et al., 2022 ^20^. The processed data for PRJCA001063 ^17^ was obtained from zenodo [10.5281/zenodo.3969339] (RRID:SCR_004129), which included cell label annotations for 10 cell types after quality control (QC) steps. Other scRNA-seq datasets were downloaded from the NCBI GEO database or as specified in their respective publications. These datasets were combined, harmonized (harmony v0.1), and prepared for downstream analysis using the bioinformatics methodology described by Chijimatsu et al., 2022 ^20^. (PMID: 31740819).

Seurat objects (RRID:SCR_022555) were created for individual datasets by reading the Cellranger output files into the R environment using the Read10x function and transforming them using the CreateSeuratObject function, as described in the Seurat Guided Clustering Tutorial (<https://satijalab.org/seurat/articles/pbmc3k_tutorial>). The analysis was conducted using R version 4.4.2 and Seurat version 5.2.01. Transcript counts, measured as UMIs, were normalized to 10,000 counts per cell and log-transformed, following the methodology described by Chijimatsu et al., 2022 ^20^. Cells with a high percentage of mitochondrial genes (>25%) were filtered out during the QC steps. Other QC metrics for individual datasets, such as UMI counts and the number of expressed genes, were also applied as per Chijimatsu et al., 2022 ^20^.

Datasets were batch-corrected and integrated using the rPCA method as outlined in the Seurat package (<https://satijalab.org/seurat/articles/integration_rpca.html>). Each dataset was scaled, and the FindVariableFeatures function was employed to identify highly variable genes. These genes were utilized for PCA analysis (RRID:SCR_014676). An anchor was created using the FindIntegrationAnchors function with the following parameters: 30 principal components, rPCA, and two reference datasets (PRJCA001063 and GSE155698). Subsequently, six datasets were integrated using the IntegrateData function of the Seurat package. The integrated dataset was scaled, followed by PCA analysis and UMAP (RRID:SCR_018217) visualization. Cell-type annotation was transferred from the reference dataset PRJCA001063 ^17^. Cellchat analyses were conducted using CellChat (v1.5.0, PMID: 39289562). The heatmaps were designed after manual selection of the topgenes and normalizing as scaling using the DoHeatmap function in Seurat (v5.2.0). The gene enrichment was performed using bitr and enrichGO from the clusterProfiler (4.14.4) and enrichplot (1.26.6) package and plotted with the ggplot2 package (3.5.2).

**mRNA-seq**

The integrity of RNA was validated by gel electrophoresis and 500ng was used to make the libraries in triplicates for each condition. Libraries for cells were made using the TruSeq RNA Library Prep Kit V3 (Illumina) according to the manufacturer’s instructions. Oligo-dT beads were used to capture poly-A tailed-mRNA followed by first-strand cDNA synthesis by Superscript II reverse transcriptase (Thermo Fischer). Second-strand synthesis was followed by end repair, 3’ adenylation, adaptor ligation, and library amplification. Agencourt AMPure XP (Beckman Coulter) was used for size selection during the library synthesis. The quality of the resulting DNA was measured with high sensitivity DNA kit (Agilent) on the Agilent TapeStation 4150. Samples were sequenced (paired-end 50bp) on a HiSeq4000 (Illumina) at the Genome Analysis Core at the Mayo Clinic (30-minute-treated RNA-seq samples) and on a NextSeq 2000 (P2, Illumina) at the Robert Bosch Center for Tumor diseases (RBCT).

**RNA-seq bioinformatic analysis**

Bam files were generated using STAR version 2.7.3a (RRID:SCR_004463) ^21^. Features were counted using htseq version 0.9.1 (RRID:SCR_005514) ^22^. Differential gene expression analysis was performed by DESeq2 (RRID:SCR_015687) ^23^. Upregulated genes were identified as ≥1 log2 Fold Change, FDR≤0.05, and BaseMean≥10. Pathway analyses were performed using ShinyGO version 0.80 (RRID:SCR_019213) ^24^.

**RNA-seq quality control and sample exclusion**

For both AsPC-1 (Fig. S3b) and L3.6pl (Fig. 3b) cell lines, one RNA-seq sample from the TNFα-treated group (replicate 1) was excluded from downstream analysis. This decision was based on pre-analysis quality control, which included evaluation of read counts, principal component analysis (PCA), and z-score heatmaps. In both cases, the excluded sample exhibited markedly low total read counts and clustered as an outlier in PCA plots. Further investigation revealed the issue was due to sequencing error. Although the exclusion criteria were not pre-established, they were applied consistently based on these objective quality metrics.

**TCGA Data Acquisition**

For RNA-seq data analysis, Gene expression data for pancreatic adenocarcinoma (PAAD) were obtained from the TCGA-PAAD project via the UCSC Xena Browser ([https://xenabrowser.net](https://xenabrowser.net/), RRID:SCR_005012). The dataset TCGA-PAAD.star_counts.tsv, containing STAR-aligned gene-level expression in log2(count + 1) format, was downloaded from the GDC Xena Hub ([https://gdc.xenahubs.net](https://gdc.xenahubs.net/)). In addition, clinical phenotype metadata were also obtained from the GDC data portal (RRID:SCR_014514) for sample annotation and filtering*.* The TCGA-PAAD dataset includes both primary tumor and normal pancreatic tissue samples and is publicly available under the accession number phs000178 in the dbGaP database (<https://www.ncbi.nlm.nih.gov/projects/gap/cgi-bin/study.cgi?study_id=phs000178>).

Sample selection was performed following rigorous preprocessing, including removal of normal tissue samples (barcode suffix -11A), tumor subtypes such as cystic, mucinous, and serous adenocarcinomas, and samples with low or missing expression of TNF and TWEAK. Gene identifiers were cleaned by removing Ensembl version suffixes and mapped to HGNC gene symbols using the biomaRt package (RRID:SCR_002987). After quality control, 82 primary tumor samples and 23,495 genes were retained for downstream analysis. Samples were then categorized into three expression groups; TNF^high^, TWEAK^high^, and TNF/TWEAK^low^, based on data-driven expression thresholds for TNF and TWEAK genes, determined using summary statistics of their expression distributions.

**TCGA Bioinformatic Analysis**

Differential gene expression was assessed using the limma package (RRID:SCR_010943) ^25^ from Bioconductor. The input matrix, already log2(count + 1) transformed, was quantile normalized using normalizeBetweenArrays() to reduce technical variability. Limma was selected over count-based methods such as DESeq2 or edgeR due to the log-transformed nature of the input data and its robust performance with small-to-moderate sample sizes. Group comparisons were defined based on TNF and TWEAK expression levels. A design matrix without intercept (~ 0 + group) was used to fit group-specific linear models with lmFit(). Empirical Bayes moderation was applied using eBayes(trend = TRUE) to account for the mean–variance relationship typical of RNA-seq data.

Genes were considered significantly differentially expressed if they met the criteria:
adjusted p-value < 0.05, |log2 fold change| > 1, and average expression ≥ 5 (log2 scale). Significant genes were saved separately for downstream visualization and enrichment analysis.

**Gene Set Enrichment Analysis (GSEA)**

GSEA was conducted using the clusterProfiler package (RRID:SCR_016884) ^26^. Gene lists ranked by log2 fold change were analyzed using the GSEA() function, which internally uses the fgsea algorithm (RRID:SCR_020938) ^27^ for fast enrichment scoring. Custom gene sets, such as TNFα_specific and Common_Genes (supplementary materials), were prepared in GMT format and loaded with read.gmt(), then passed via the TERM2GENE argument. The analysis used the following parameters; pvalueCutoff = 1, minGSSize = 1, maxGSSize = 600. Documentation for the GSEA function is available in the clusterProfiler user manual ^28^. Enrichment statistics, including normalized enrichment score (NES), nominal p-values, and leading-edge genes, were computed. Selected gene sets were visualized using the plotEnrichment() function, and results were saved in TSV and PDF formats for reporting.

Analyses were conducted in R v4.3.1 using the following packages: limma v3.60.6, clusterProfiler v4.12.6, fgsea v1.30.0, and biomaRt v2.60.1.

**Chromatin immunoprecipitation sequencing (ChIP-seq)**

Chromatin immunoprecipitation was performed as previously described ^3,29^. Antibodies included H3K27ac (1μg; C15410196, Diagenode; RRID:AB_2637079), FOSL1 (D80B4) (FRA-1) (5 μl; 5281; Cell Signaling; RRID:AB_10557418), RELA (NFκB p65 L8F6) (5 μl; 6956; Cell Signaling; RRID:AB_10828935), RELB (5 μl; Cell Signaling; 10544; RRID:AB_2797727), and control Rabbit IgG (1 μg; C15410206, Diagenode; RRID:AB_2722554). Protein A-sepharose and protein G-Sepharose (for RELA) beads were added to the samples and incubated for 2 hours, washed, de-crosslinked, and DNA was extracted. Samples were performed in triplicates for each. Condition. Libraries were prepared using the MicroPlex Library Preparation Kit v2 (Diagenode) according to the manufacturer’s protocol. DNA integrity was measured with a high-sensitivity DNA kit (Agilent) on the Agilent TapeStation 4150. Samples were sequenced (paired-end 50bp) on a NextSeq 2000 (P3, Illumina) at the Robert Bosch Center for Tumor diseases.

**ChIP-seq bioinformatic analysis**

Reads were mapped to the reference genome assembly (hg38) by BOWTIE2/2.5.0 (RRID:SCR_016368) ^30^. Bigwig files were generated from merged bam files using bamCoverage (RRID:SCR_016366). Localization profiles were viewed using Integrative Genomics Viewer (IGV 2.16.0) (RRID:SCR_011793) ^31^. MACS2 (RRID:SCR_013291) ^32^ was used to call the significant peaks without building the shifting model with broad peaks (broad-cutoff 0.05) called for H3K27ac, narrow peaks (broad-cutoff 0.05), and input files from respective cells as background. The Bioconductor (RRID:SCR_006442) R package Diffbind 3.10.1 (RRID:SCR_012918) ^33^ was run on R version 4.3.1 according to the instruction manual to define regions that are differentially enriched by RELA and RELB. ChIP occupancies were evaluated by the computeMatrix tool (RRID:SCR_016366). ChIP-seq profiles and heatmaps were generated from computeMatrix values using the PlotProfiles and PlotHeatmap tools respectively on the Galaxy platform (RRID:SCR_006281) ^34^. Transcription factor enrichment analyses were performed using ChIP-Atlas (RRID:SCR_015511) ^35^.

**Multiplex immunofluorescence staining**

A six-color multiplex immunofluorescence staining was performed using OPAL^TM^ multiplexing method. The staining protocol for FFPE tissue sections was optimized for the simultaneous detection of six antibodies and DAPI for cell nuclear stain. The sections were deparaffinized, rehydrated, subjected to heat-induced epitope retrieval, and incubated with primary and secondary antibodies. The antibodies were visualized using a fluorescent tyramide with Opal 6-Plex Manual Detection Kit (Akoya Biosciences; NEL861001KT). The epitope retrieval and staining process was repeated sequentially for different primary antibodies and fluorescent tyramide combinations. The following primary antibodies with different dilutions were used: CD68 (Cell Marque; 168M-94; RRID:AB_1158188) with 1:500 dilution, Phospho-NF-kB p65 (Ser536) (Cell Signaling; 3033; RRID:AB_331284) with 1:25 dilution, αSMA (Abcam; ab5694; RRID:AB_2223021) with 1:100 dilution, CD31 (PECAM-1) (89C2) (Cell Signaling; 3528; RRID:AB_2223021) with 1:800 dilution, and Pan-Keratin (AE1/AE3) (Cell Signaling; 67306) with 1:50 dilution. Antibodies were visualized with the following tyramide dyes used from the Opal Detection kit (Akoya Biosciences; NEL861001KT): Opal 520, Opal 570, Opal 620, Opal 690, and DIG-Opal 780. Sections were mounted with ProLong^®^ Diamond Antifade Mountant (Thermo Fischer Scientific; P36961). Multiplex-stained slides were imaged using a PhenoImager Fusion system (Akoya Biosciences).

35.	Oki SO, T. ChIP-Atlas. <https://chip-atlas.org>
