## Supplementary figures and images for "Epigenetic Context Defines the Transcriptional Activity of Canonical and Noncanonical NF-κB Signaling in Pancreatic Cancer"

### Supplemental Material 2

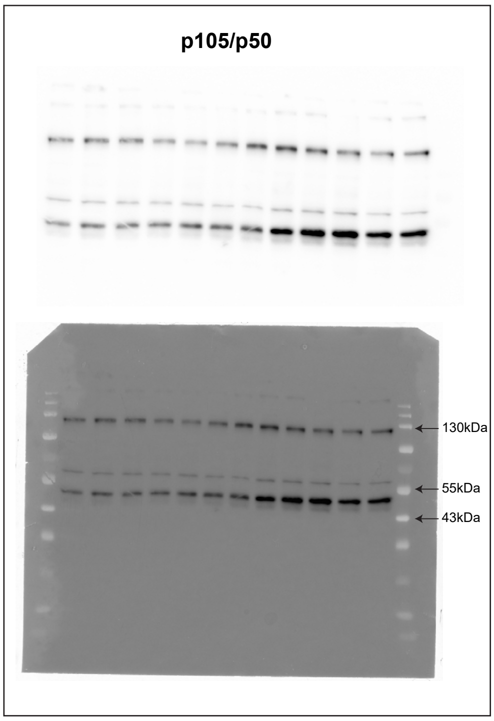

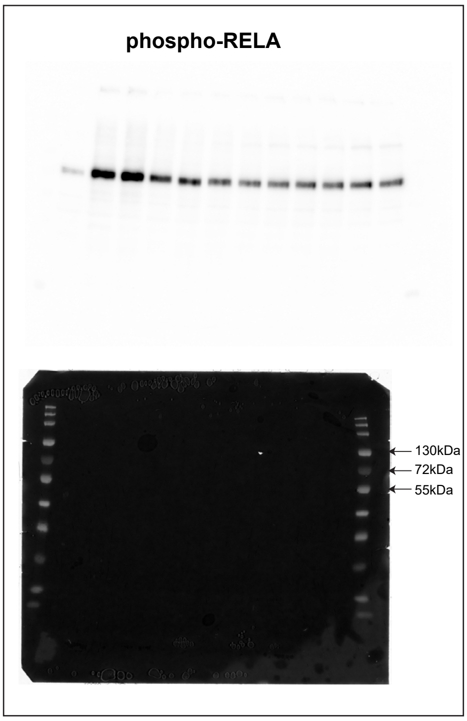

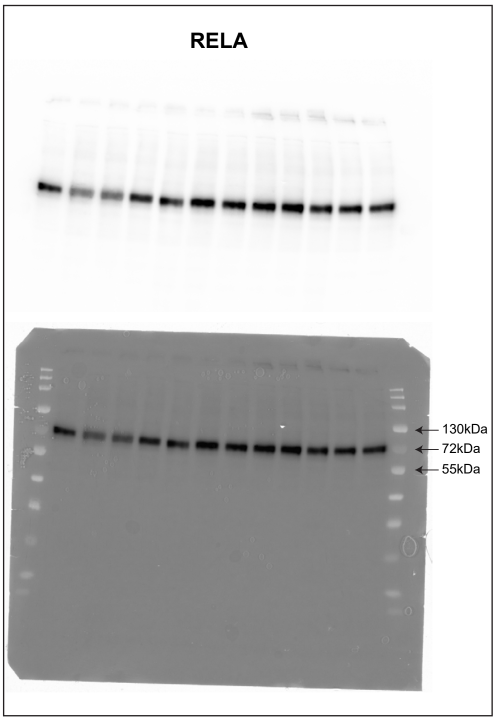

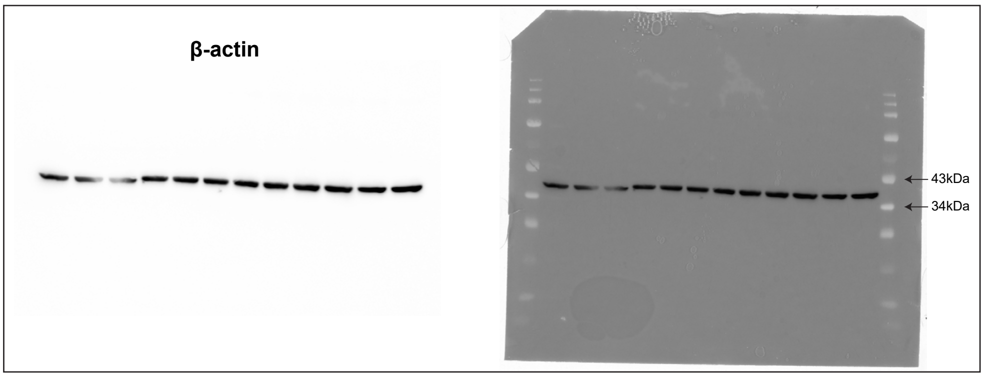

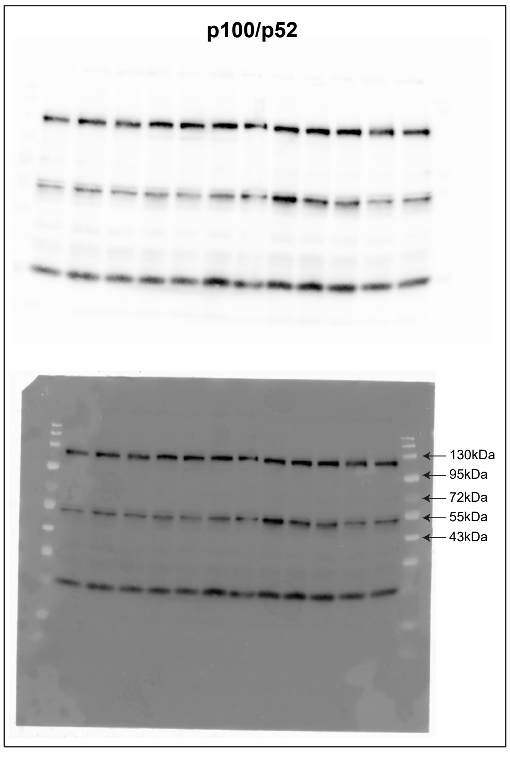

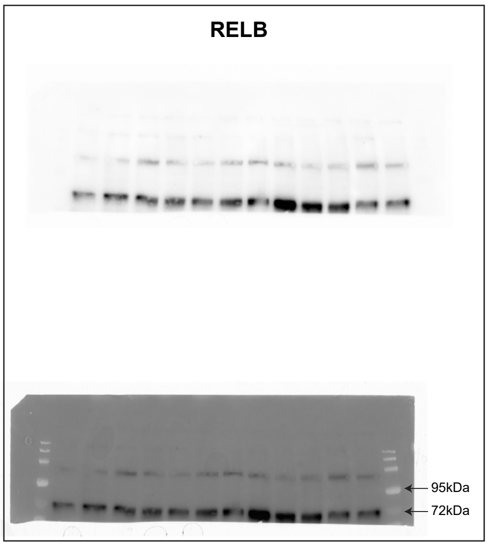

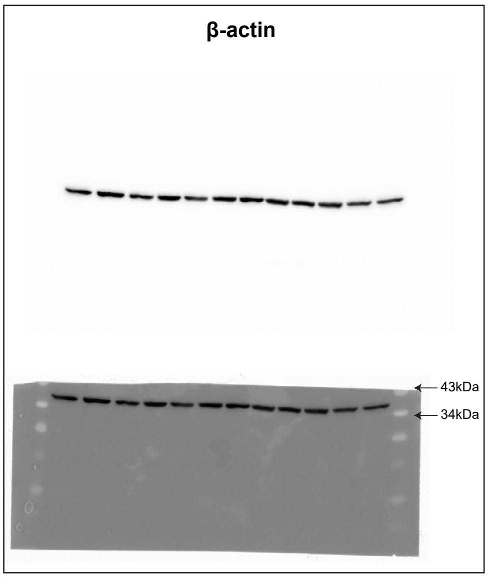
