## Supplementary Figures S1-S5 for "Epigenetic Context Defines the Transcriptional Activity of Canonical and Noncanonical NF-κB Signaling in Pancreatic Cancer"

#### Supplementary Figure 1

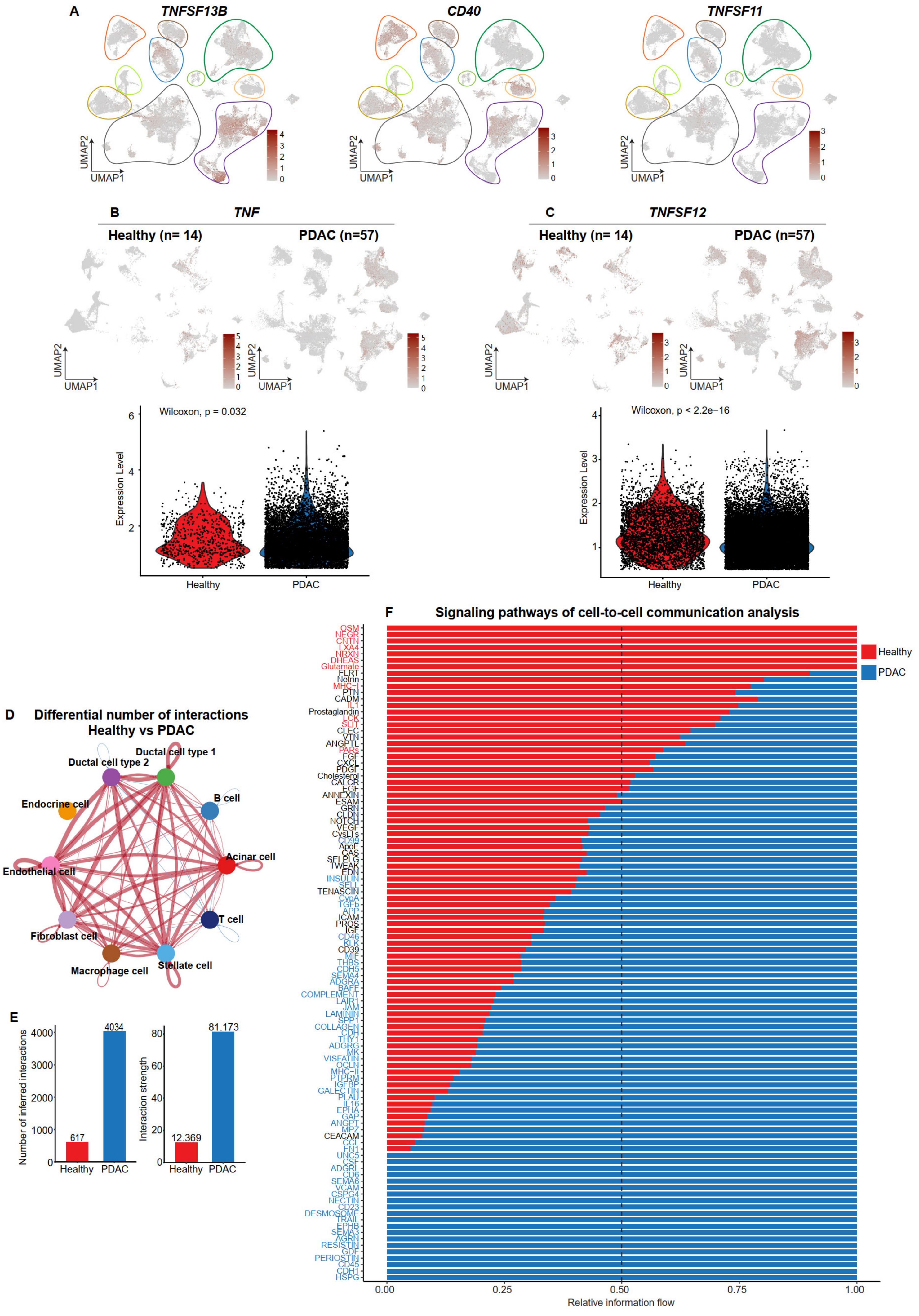

Supplementary Figure 2

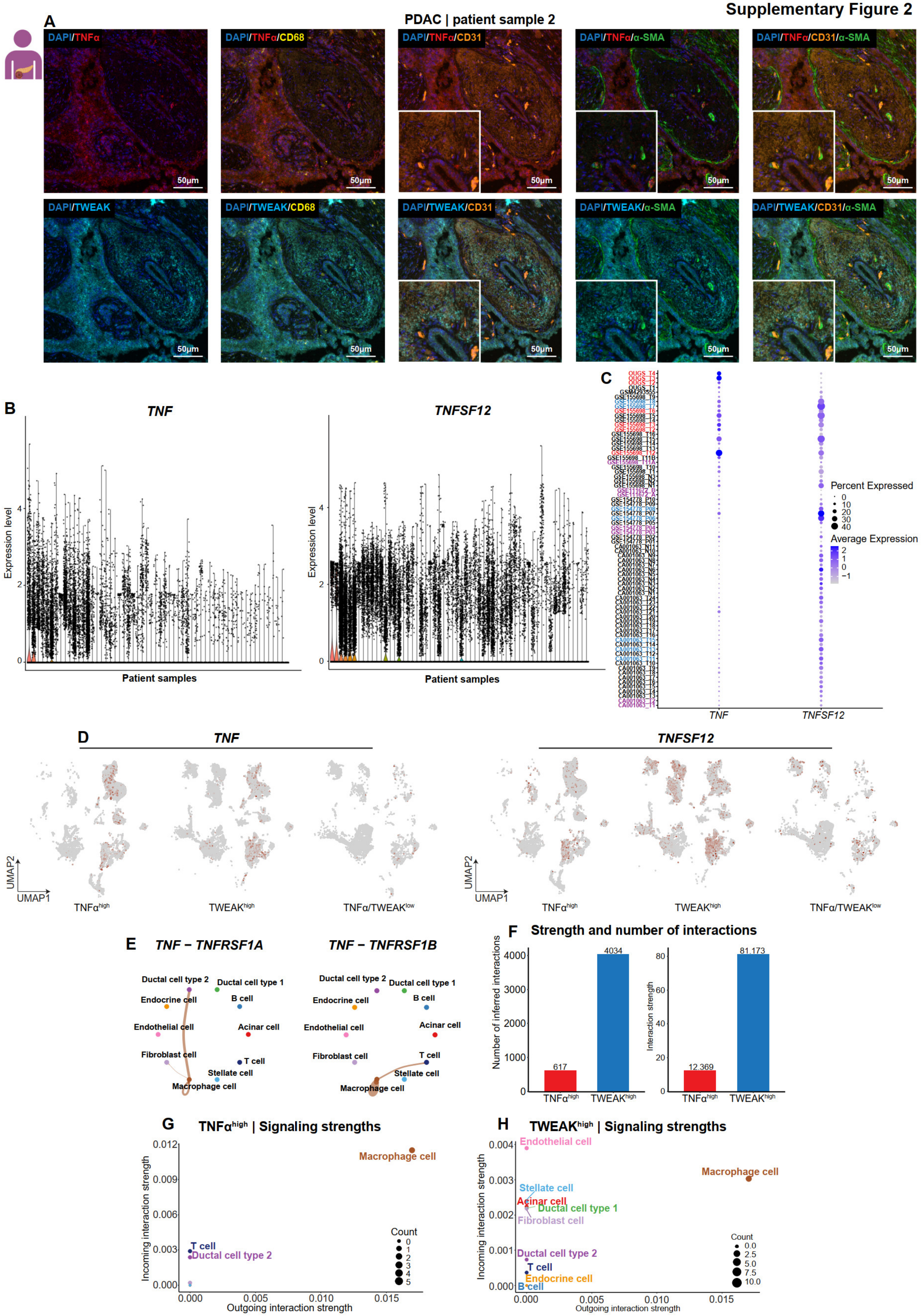

#### Supplementary Figure 3

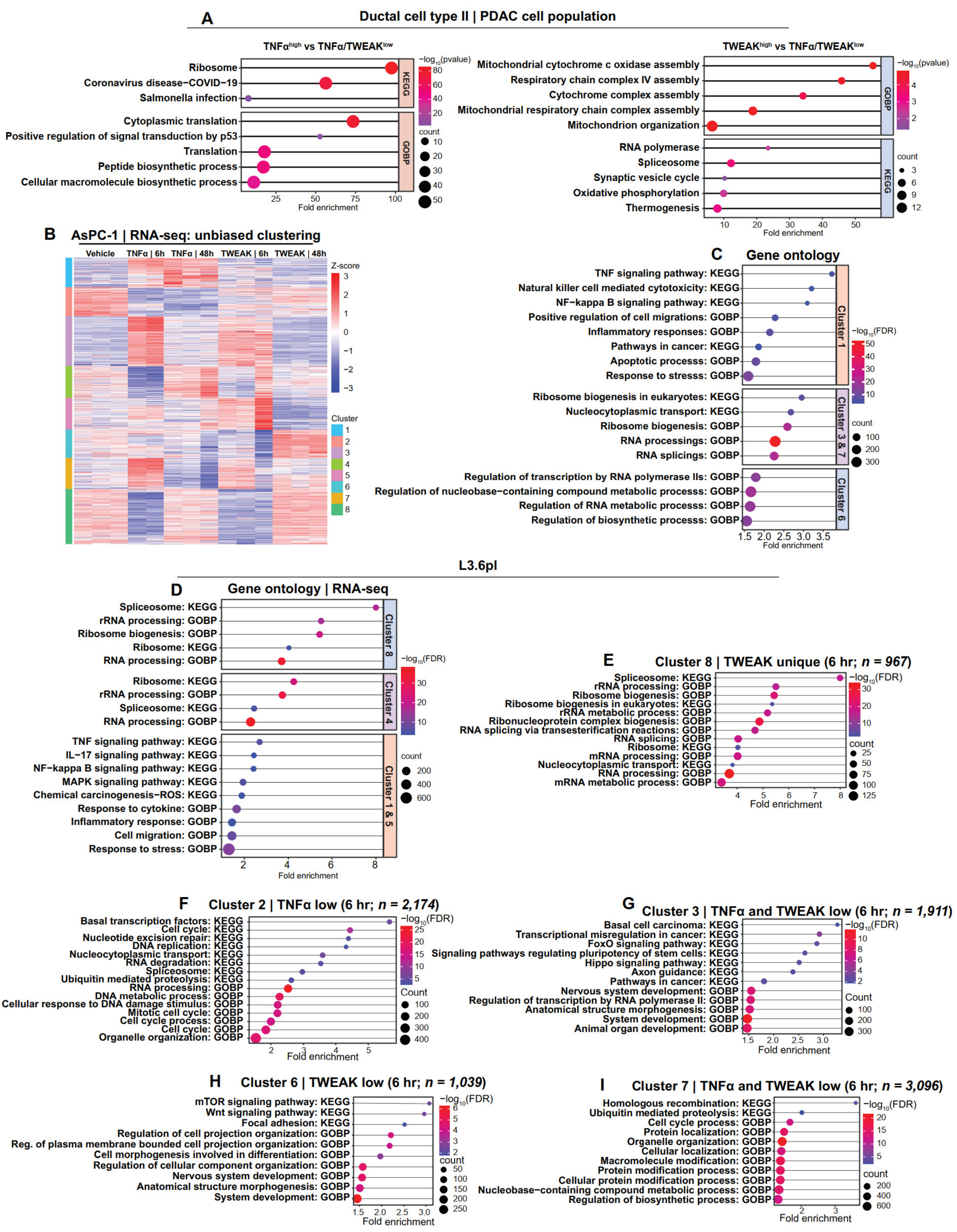

### Supplementary Figure 4

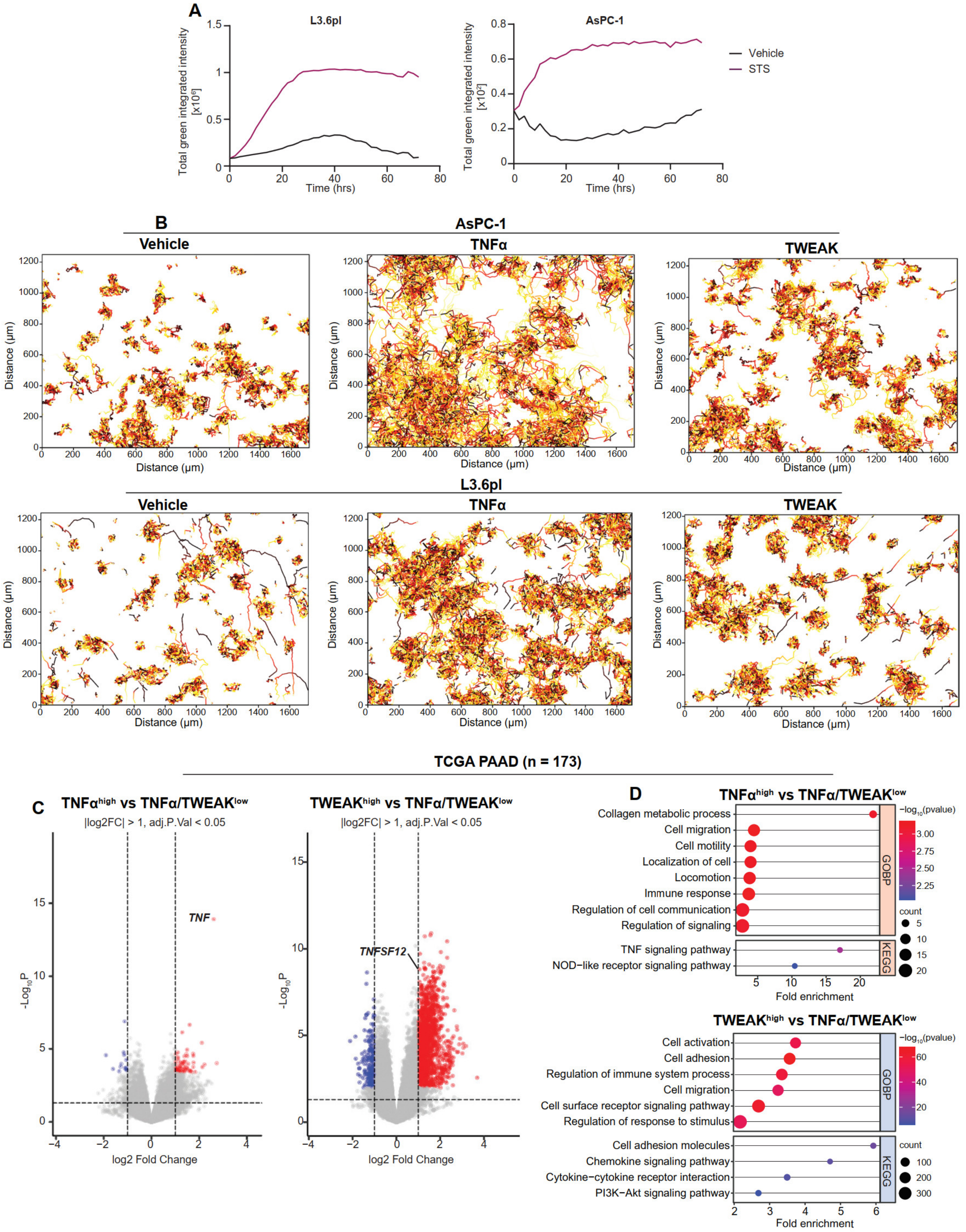

**Supplementary Figure 5**

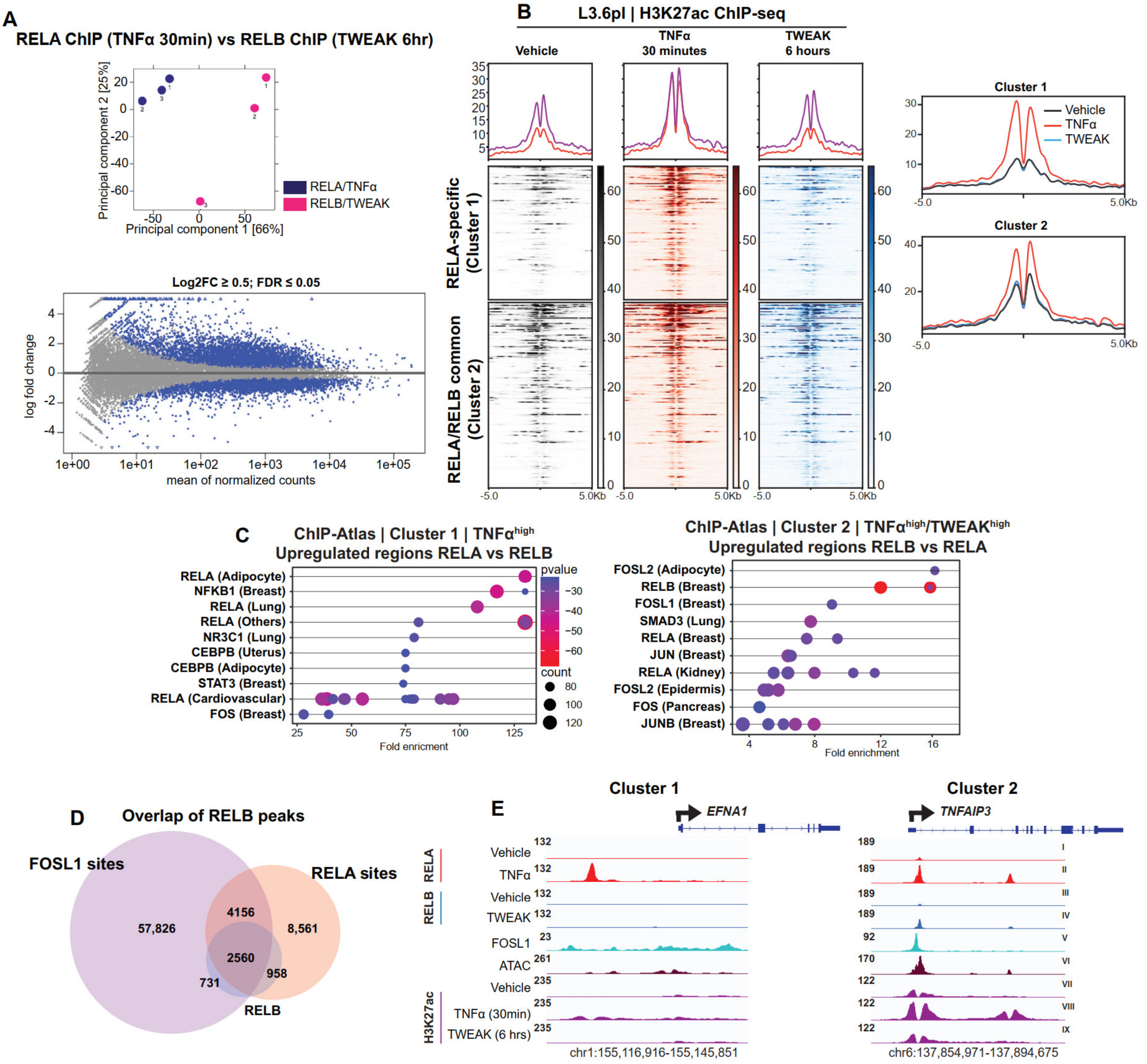
